# Time-dependent BMP4 signaling directs lineage specification in human mesoderm

**DOI:** 10.64898/2026.07.03.736254

**Authors:** Wei Zhao, Filip J. Wymeersch, Minoru Takasato

## Abstract

Human pluripotent stem cells (hPSCs) provide a powerful platform for modeling early human embryonic development. Here, we investigate the mechanisms underlying mesodermal heterogeneity using a minimal directed differentiation system that simultaneously generates paraxial (PXM), intermediate (IM) and lateral plate mesoderm (LPM) populations. Single-cell RNA sequencing across defined time points during hPSC differentiation revealed a temporal sequence of lineage specification with LPM emerging first, followed by PXM and IM differentiation. Ligand-receptor and differential gene expression analyses identified BMP4 as a key regulator enriched in LPM-associated clusters versus mesoderm progenitors (MPs) that hold PXM and IM precursors. Whereas LPM cells cluster with an early BMP4 signal, IM clusters are associated with later BMP4. Moreover, these early and late BMP4 signals regulate this lineage specification potentially through distinct downstream pathways. Leveraging this insight, we established a stepwise protocol combining early BMP inhibition with subsequent BMP4 supplementation, suppressing initial LPM fate to efficiently induce IM from a mixed MP population. Longer culture of these selective IM progenitors promotes more mature nephrogenesis. Moreover, we demonstrate that during early differentiation high levels of BMP4 can still redirect MPs to more lateroventral fates, illustrating a degree of plasticity within the mesoderm lineage. Together, our results define a temporal framework for BMP4 signaling in mesoderm fate determination and provide a strategy for selective mesoderm differentiation from hPSCs.

**HIGHLIGHTS:**

- Development of a minimal 2D differentiation platform allows for heterogenous mesoderm formation.
- Temporal BMP4 signaling differentially directs mesoderm fates, with early exposure favoring LPM and late exposure promoting IM identity.
- LPM cells arise first while later mesoderm progenitors hold both IM and PXM-fated cells.
- Sequential BMP modulation promotes IM and enhances nephrogenesis.

## INTRODUCTION

During gastrulation, the mammalian embryo generates different types of mesenchyme from the primitive streak (PS), including the paraxial, intermediate and lateral plate mesoderm (PXM, IM and LPM), that will go on to form distinct tissues from the dermis and musculoskeletal system^1,2^, the urogenital organs^3–5^, and heart, limbs, genitalia and other visceral structures^6,7^, respectively. Single cell and small population fate mapping during mouse and chick embryo development have shown different mesoderm populations arise along the length of the PS at different stages of gastrulation^8,9^ (reviewed by Tam and Loebel, 2007^10^). The pattern of mesoderm fate distribution in the PS changes over time as certain mesoderm precursors exit through the streak, *e*.*g*. cardiac or extra-embryonic mesoderm progenitors, while other mesoderm progenitors remain in the epiblast. In mid-to-late streak-stage mouse embryos, LPM cells arise from the mid-posterior PS, whereas PXM cells arise from the anterior PS^8,11^. Later, at early somitogenesis, neuromesodermal progenitors in the caudal embryo give rise to somite and spinal cord tissues, whereas the posterior-most PS fraction still contains lateral mesoderm progenitors that will form the caudal cloacal mesenchyme^12–14^, suggesting new LPM formation has concluded by early somitogenesis stage. The exact developmental origin of the IM, however, is less defined as fate mapping in mouse and chick embryos show only few progenitors can be labelled directly and IM-fated cells are often found alongside co-labelled descendants in the LPM or PXM, both in gastrulation and early-somite stage embryos^15,16^. Moreover, lineage-tracing and tissue grafting have shown the type of IM progenitor produced changes during this period from ‘anterior’ to ‘posterior’ IM (aIM and pIM), creating the Wolffian duct and metanephric mesenchyme, respectively^5,13,17^. Interestingly, despite the divergent axial tissues they will eventually give rise to, cells leaving the caudal progenitor zone can still be redirected to another mesoderm fate either by providing them with different signaling cues^12,18^ or modulating fate-committing transcription factors^19^, suggesting a degree of plasticity within the mesoderm. As gastrulation and axial elongation proceed, the fate of these three main mesoderm lineages will become fixed, however, the timing as well as the drivers of this plasticity in PXM/IM/LPM progenitor populations remains less studied. Moreover, how mesoderm lineage commitment proceeds over time during early embryo development remains unclear, especially during human embryogenesis.

Whereas the genetic mechanisms in mesoderm patterning have been well characterized in model organisms, those that govern lineage specification during *in vivo* human development remain less understood due to ethical/legal restrictions and the rarity of early gestation human embryonic material^20^. Human embryonic and induced pluripotent stem cells (hESCs/hiPSCs) provide an alternative, tractable system to model early human development *in vitro*, and various protocols have shown differentiation towards PXM^21–23^, IM^24–26^ and LPM-derived tissues and organoids^22,27–29^. Many of these protocols first use a canonical WNT signaling to induce PS-like mesoderm populations and further diverse strategies, including WNT, Nodal/TGFβ, FGF and BMP pathways modulation, to direct cells to a distinct anteroposterior PS identity. In general, it has been shown that BMP signaling regulates early lineage bifurcation between PXM and LPM^22,23,30^, however, the signaling cues that direct cells to the IM lineage are less well defined. This has resulted in multiple differentiation strategies that successfully generate IM *in vitro*^32^. Previously, we have used high concentrations of CHIR99021, a GSK3β inhibitor and WNT signaling agonist (CHIR), to direct hiPSCs towards a posterior PS intermediate. Other studies have used both BMP addition^17,24,31–33^ as well as BMP pathway inhibitors like Noggin^25^ or LDN193189^24^ during IM differentiation to obtain nephrogenic cell derivatives^34^. Moreover, some protocols additionally stimulate the Nodal/TGFβ pathway^17,24,25,33^, often associated with more anterior PS fates^10^, followed by a posteriorizing BMP4 signal to achieve successful IM induction^17,33^. Taken together, this diversity in successful IM protocols likely reflects both the plasticity in the mesoderm and the timed derivation of aIM and pIM fates *in vivo*, as well as the likelihood of a mixed mesodermal origin of the nephrogenic mesenchyme^35^. BMP4 has been shown to control lineage specification along the mediolateral axis *in vivo*^36–38^ and steer LPM-PXM fate during *in vitro* PSC differentiation^22^. However, how BMP4 - or other signaling pathways - regulates IM specification remains poorly defined.

To elucidate the mechanisms governing mediolateral fate decisions during human mesoderm differentiation, we established a minimal 2D directed system of hiPSCs differentiation that enables the simultaneous induction of heterogeneous mesoderm populations, including PXM, IM and LPM. Using single-cell RNA sequencing (scRNA-seq) across defined time points, we identify a temporal transcriptional hierarchy in mesoderm specification and key signaling dynamics underlying lineage divergence. Furthermore, using timed BMP modulation experiments, we further investigated the developmental competence and plasticity of MPs. Guided by these approaches, we define a temporal framework for BMP4 signaling in human mesoderm patterning, we provide a novel, stepwise protocol for IM derivation and subsequent maturation of the nephrogenic mesenchyme.

## RESULTS

### Generation of heterogeneous mesoderm populations from hiPSCs

To model mesoderm diversification *in vitro*, we established a differentiation system that enables the simultaneous induction of heterogeneous mesoderm populations from hiPSCs with minimal factors (**Figure 1A-B**). We first optimized the cell culture conditions. Cell density is known to critically influence mesoderm lineage composition, with lower seeding densities favoring *PAX3*^+^ PXM differentiation^39^ and higher densities promoting *FOXF1*^+^ LPM identity^40^. We found that seeding 3.5 × 10^4^ hiPSCs/cm^2^ on day (D)0 of differentiation was most optimal to simultaneously induce PXM, IM and LPM (**Figure S1A-B**). Titrating the CHIR concentration showed that high CHIR levels (8 μM) preferentially induce *PAX2*^+^ IM differentiation^41^ while concomitantly suppressing *PAX3*^+^ PXM and *FOXF1*^+^ LPM cells, reminiscent of our published kidney organoid protocol^26^. Lower CHIR concentrations (≤5 μM) supported co-induction of PXM and LPM, with further reduction to 3 μM leading neural lineage marker expression, including *SOX2* and *FOXD3*, indicating a potential loss of mesodermal identity (Figure S1C). Based on these results, we seeded the cells at 3.5 × 10^4^ hiPSCs/cm^2^ and activated WNT signaling with 4µM CHIR for three days, followed by six days in basal medium (APEL2 with 3% PFHM-II).

**Figure 1.**
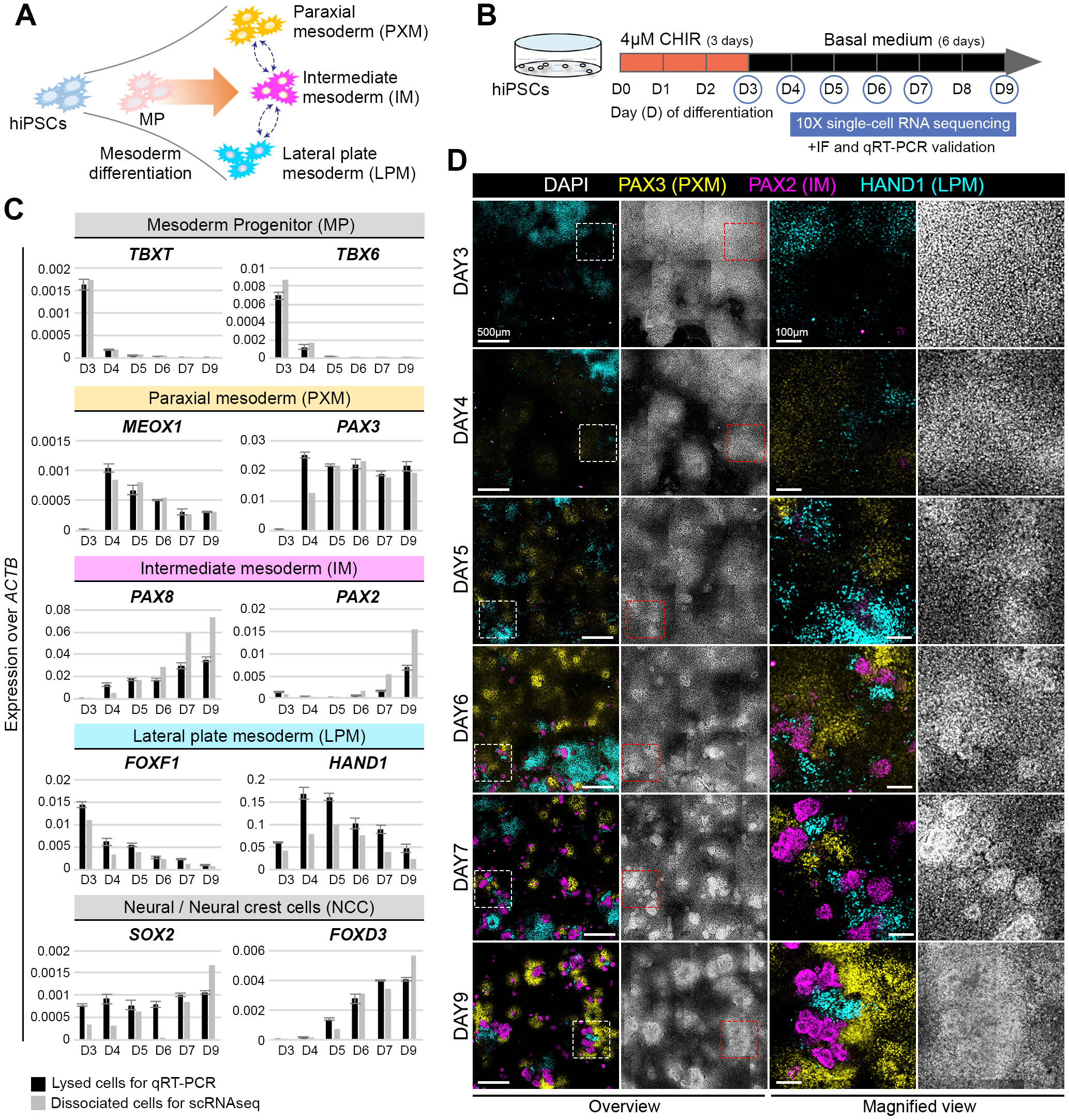
Induction of heterogeneous human mesoderm. **(A)** Schematic of hiPSCs differentiation towards multiple mesodermal lineages (orange arrow) and plasticity between them (dashed blue arrows). **(B)** Nine-day differentiation protocol with 4 μM CHIR99021 (CHIR) for 72h, followed by culture in basal medium until D9. **(C)** qRT-PCR analysis of lineage markers in induced cells for qRT-PCR (grey) and in samples used for scRNA-seq (black). **(D)** Immunofluorescent staining at different days against PAX3 (PXM), PAX2 (IM) and HAND1 (LPM), counterstained for nuclei with DAPI. Images are stitched orthogonal projections of confocal stacks of one well (48-well plate).

We then performed quantitative reverse transcription polymerase chain reaction (qRT-PCR) on D3-D7 and on D9 of differentiation to assess mesoderm progenitor (MP), PXM, IM and LPM expression marker dynamics (Figure 1C). MP markers *TBXT* and *TBX6* were downregulated by D4 and largely absent by D5. Three mesoderm types were timely generated, shown by their well-described marker expression^22,26^: *MEOX1* and *PAX3* for PXM, *PAX8* and *PAX2* for IM, and *FOXF1* and *HAND1* for LPM. *SOX*2^+^ neural, *FOXD3*^+^ neural crest cells (NCCs; Figure 1C) or *SOX17*^+^ endothelium cells were minimally induced in our protocol (Figure S2A-B). Immunofluorescent staining further confirmed the coexistence of multiple mesodermal lineages in our protocol and from D6 onward, characterized by simultaneous, non-overlapping PAX3, PAX2 and HAND1 expression (**Figure 1D**). At later days, cells expressed markers associated with somitogenesis, such as *UNCX, TCF15* and *TBX18* as well as nephrogenesis markers, including *SIX2, LHX1* and *GATA3*, suggesting nephron progenitor^42,43^, early epithelial structures^44^ and distal tubular lineages^45^ had formed. By D9 of differentiation, nephron-like structures containing multiple segment identities could be clearly observed (**Figure S2B-C**), suggesting this protocol can support further nephrogenesis.

We then performed scRNA-seq on harvested cells post-CHIR withdrawal on D3-D7 and on D9 of differentiation and compared them to the obtained qRT-PCR results to assess whether single cell dissociation had introduced any bias in mesoderm proportions (**Figure 1B-C**). Most lineage markers showed comparable expression levels between the two conditions, indicating efficient and representative recovery of mesodermal populations. Epithelial-associated markers *PAX2*, PAX8 and *LHX1* were slightly enriched following dissociation, likely reflecting preferential recovery of cells in organoid-like structures. In contrast, LPM markers *FOXF1* and *HAND1* were modestly reduced, possibly due to stronger adhesion to the culture substrate (Figure 1C and S2B). Taken together, these results show we established a robust *in vitro* system that recapitulates heterogeneous mesoderm differentiation with minimal initial intervention.

### Temporal dynamics of mesoderm lineage specification at the single cell level

We went on to capture the differentiation dynamics in our system at the single cell level. In total 13,967 cells were recovered over six days, grouping in 16 distinct cell populations by unsupervised clustering and annotated based on established lineage markers (**Figure 2A-C and S3**). Endothelial and neural populations were detected at low frequency and confined to minor clusters (*10* and *13*). Only two small clusters (*11* and *14*) could not be identified by specific markers, suggesting limited off-target differentiation occurred in our protocol (**Figure 2C, S3** and **S4A-B**). Although D3 cells largely exhibited a MP-like identity, marked by *TBX6, MIXL1* and *CDX2* expression (*cluster 2*), a subset of cells already expressed LPM markers, including *FOXF1* and *HAND1*, indicating the early emergence of LPM. *HAND1* expression remained in this LPM population until D9 and thus formed a good marker for LPM fate (*cluster 1*; **Figure 2C-D**). We also found cardiac mesoderm progenitor and endothelium markers in this population, suggesting an early LPM identity (Figure S4B-C). On D4, three major clusters emerged (*cluster 3, 8* and *9*) which we identified as the LPM, a population of mixed mesoderm and cell cycle gene-expressing cells (LPM d4, MM d4 and CC, respectively; Figure 2B-D). At this time, PXM markers *PAX3* and *MEOX1* became expressed, accompanied by the early IM marker *OSR1* (**Figure 2C-D**). However, cells did not diverge at this point, suggesting somitic and more lateral fates including the IM are linked as suggested by clonal labelling and lineage reconstructions in mouse embryos^46,47^.

**Figure 2.**
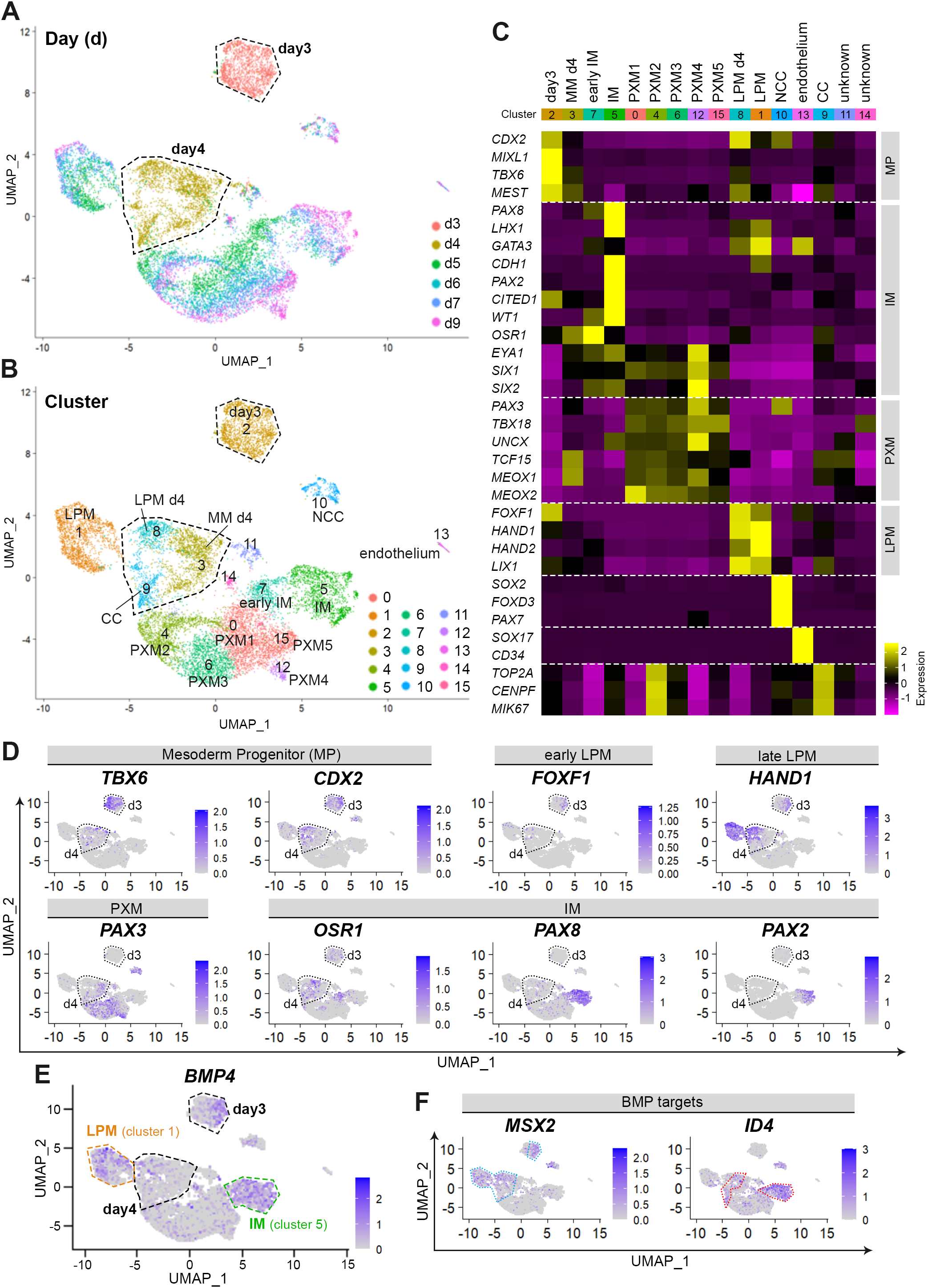
Clustering analysis shows mesoderm lineage specification at the single cell level. **(A-B)** Uniform Manifold Approximation and Projection (UMAP) plots showing single cells across days of differentiation **(A)** and cluster **(B). (C)** Heatmap of cluster-specific marker gene expression. **(D-F)** UMAP feature plots showing mesoderm progenitor, mesoderm lineage specification markers **(D)** and *BMP4* expression **(E)** with downstream targets, *MSX2* and *ID4*, across all clusters **(F)**. Dashed lines in A, B, D, E, approximate location of D3 and D4 cells. Orange and green dashed lines in E, approximate location of LPM and IM clusters, respectively. Blue and red dotted lines in F, approximate location of *MSX2*^+^ and *ID4*^+^ cells.

D5-7 and D9 cells showed further separation of mesoderm fates. Initial parallel emergence of the PXM and IM lineages, diverged as IM maturation progressed with increasing *PAX8* expression from D4 and *PAX2* expression from D6 which remained largely restricted to IM-associated clusters (*cluster 5* and *7*). Five PXM-like cell groups clustered together expressing *TCF15, TBX18, UNCX, MEOX1, MEOX2* and *PAX3* (*cluster 0, 4, 6, 12* and *15*), with the highest PXM-associated gene expression in the most distant *cluster 12* (**Figure 2B-C** and **S4D**). Interestingly, IM *clusters 5* and *7* expressed higher markers for nephric tubular fate such as *LHX1, CDH1* and *GATA3*^26^, whereas PXM and IM clusters both expressed genes normally associated with nephric mesenchymal fate, such as *EYA1, SIX1* and *SIX2*^26^, likely due to its shared progenitor on D4 and reflecting the timed emergence of these kidney lineages *in vivo*^26^ (**Figure 2C-D**). Analysis of kidney-associated gene expression further supported progressive nephrogenic differentiation with nephron progenitor markers *SIX1/2* and *CITED1* and stromal lineage marker *FOXD1* detected from D5 onward, followed by enrichment of epithelial markers *LHX1, PAX2, GATA3* and *CDH1* at later stages (**Figure S4E**). This sequential activation is consistent with developmental progression of the nephric lineage^4^.

Since BMP signaling drives PXM-LPM lineage bifurcation^22,37,38,48^, we looked at *BMP4* expression and observed activation at D3, reduced expression at D4 and re-activation in *cluster 1* (LPM) and *cluster 5* (IM) from D5-D9 (**Figure 2E**). We looked at various BMP receptors and downstream targets (**Figure S5**) and noted that *MSX2* and *ID4* clustered remarkably well with LPM and IM fates respectively, suggesting a potential different downstream response to BMP4 in these two populations (**Figure 2F**). Thus, this temporal transcriptional profile analysis in a minimal intervention mesenchyme differentiation protocol reveals a clear hierarchy in mesoderm lineage specification, with LPM emerging earliest, followed by PXM and early IM, with subsequent maturation of these three mesoderm lineages.

### BMP4 is an early regulator in mesoderm lineage specification

As we observed an asymmetrical distribution of LPM markers as well as *BMP4* expression and its targets in the overall clustering at D3 and D4 of differentiation, we went on to perform sub-clustering of these days (Figure 3A). On D3, cells clustered into six groups. Marker-based annotation revealed that *BMP4* as well as *MSX2* were highly expressed in CDX2^high^ TBX6^low^ cells, alongside LPM markers *FOXF1* and *HAND1*, suggesting these cells are of lateral fate (*LPM cluster*). Another cluster showed lower *BMP4*/*MSX2/FOXF1/HAND1* expression, suggesting it might be a precursor population of the former (*pre*.*LPM cluster*). The other four clusters were either expressing higher levels of cell cycle genes (*CC1/CC2 clusters*) or expressed high levels of *TBX6* with lower *CDX2*, suggesting it might represent a nascent/mixed mesoderm population (*MM1/MM2 clusters*; **Figure 3B-D**). Expression of *HAND1*^+^ LPM cells showed a consistent overlap with *BMP4* and *MSX2*-expressing cells (**Figure 3E** and **S6B**). Next, we performed differentially expressed genes (DEGs) and ligand-receptor analyses of the *HAND1*^+^ LPM cells versus the *FOXF1*^-^ *HAND1*^-^ population to identify other potential signaling governing early fate diversion. This brought BMP4 as the top candidate regulator enriched in LPM-associated clusters with 1.5-fold change versus the *FOXF1*^-^ *HAND1*^-^ population. Serine Protease 23 (*PRSS23*) was also significantly upregulated and has been shown to be involved in EMT of the vasculature^49^. WNT signaling antagonist *DKK1* as well *FGF8/17* signaling were also significant but presented a low fold change compared to *HAND1*^+^ LPM cells (**Figure 3F, S6A** and **Supplemental Data 1**).

**Figure 3.**
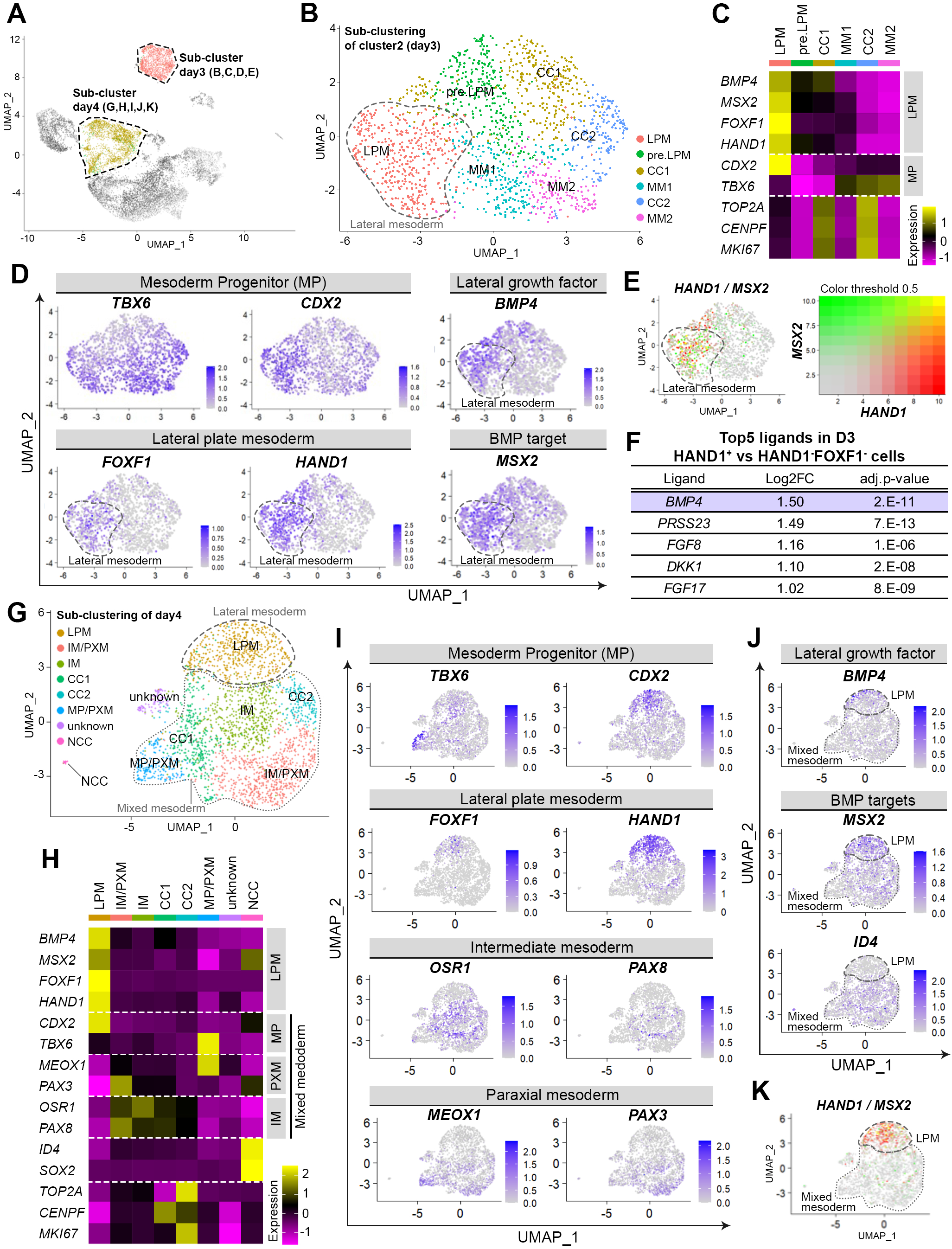
Sub-clustering shows BMP4 is an early regulator in mesoderm lineage specification. **(A)** Approximate location of D3 and D4 cells in the hiPSC differentiation system UMAP plot. **(B)** UMAP plot showing six sub-clusters in D3 cells. **(C)** Heatmap showing D3 cluster-specific marker gene expression. **(D)** D3 UMAP feature plots showing single maker expression for mesoderm progenitors (MP), LPM, and *BMP4* and *MSX2*. **(E)** Feature plot showing co-expression of *HAND1* and *MSX2* on D3. Dashed line in B, D and E, approximate location of D3 lateral mesoderm. **(F)** Top five ligands in *HAND1*^+^ LPM versus *HAND1*^-^ *FOXF1*^-^ cells on D3. **(G-H)** UMAP plot showing eight sub-clusters on D4, with heatmap showing cluster-specific marker gene expression. **(I-J)** UMAP feature plots in D4 sub-clustered cells showing MP and mesoderm lineage specification markers, *BMP4* and its downstream targets, *MSX2* and *ID4*. **(K)** Feature plot showing co-expression of *HAND1* and *MSX2* on D4. In insets G, J and K, dashed lines indicate the approximate LPM, and the dotted line, mixed mesoderm progenitors.

In D4 sub-clustering (**Figure 3G**) we identified eight clusters. Marker-based annotation showed one unknown cell cluster and NCCs (**Figure S6E**), as well as further distinction in the mesoderm with a *CDX2*^*high*^ *TBX6*^*low*^ *FOXF1*^+^ *HAND1*^+^ LPM population (*LPM cluster*) and a *TBX6*^+^ *MEOX*^+^ population (*MP/PXM cluster*). Four other clusters showed either high levels of cell cycling genes (*CC1/CC2 clusters*) or presented markers for both IM and PXM fate, such as *FOXC1/2, PAX3, PAX8* and *OSR1* (*IM* and *IM/PXM clusters*). Interestingly, both *CC1* and *CC2* clusters also expressed low levels of *PAX3, PAX8* or *OSR1*. Together with their close UMAP clustering, it suggests all four clusters likely represent a mixed paraxial-intermediate mesoderm progenitor population (**Figure 3G-I** and **S6D**). At D4, *BMP4* was reduced and potentially reflected changes in WNT signaling following removal of CHIR (**Figure 3J**). Notably, however, *MSX2* remained specifically associated with *FOXF1*^+^*HAND1*^+^ LPM population (**Figure 3K** and **S6C**). Although IM and PXM populations showed partial overlap at the cluster level, lineage-specific marker expression was largely segregated at the single-cell level, indicating that lineage divergence has already started (**Figure 3I**). Interestingly, DEG analysis in D4 clustering identified *ID4*, a downstream target of BMP signaling, enriched in *OSR1*^+^ */ PAX8*^+^ IM-associated populations. In the overall clustering, it was also specifically enriched in IM-biased cells, suggesting it might have role in IM maturation, alone or in association with other *ID* genes (**Figure 2D-F, 3I-J** and **S5B**). Consistent with the role for BMP signaling in LPM specification at D3, *HAND1*^+^ cells overlapped well with *MSX2* at D4 (**Figure 3K**). To further assess whether these lineage-associated transcriptional BMP patterns are conserved *in vivo*, we examined single-cell and spatial transcriptomic atlases of mouse and human embryos^20,50–52^. In both species, *MSX2* expression was enriched in *FOXF1*^+^ *HAND1*^+^ LPM-like mesoderm, whereas *ID4* expression overlapped with *PAX2*^+^ *and PAX8*^+^ in mesonephric cells during later human developmental stages. (**Figure S7**).

Taken together, D3/4 sub-clustering shows LPM cells arise first from a subset of progenitors with elevated *BMP4/MSX2* signaling as the main bifurcation signal, reflecting lateral mesoderm formation *in vivo* and *in vitro*^36–38,48,53^. PXM and IM lineages cluster later and together at D4, but are distinct at the single cell level, suggesting they likely share a common mesoderm precursor for longer. Moreover, early signs of PXM-IM bifurcation appear to be *BMP4*/*ID4*-driven.

### BMP4 exerts temporally distinct effects on LPM and IM specification

To test these temporal effects of BMP4 on mesoderm lineage specification, we added 5 ng/mL BMP4 for 24 h at different days of differentiation ([D2-3], early; [D3-4], mid; or [D4-5], late BMP4 exposure) and performed qRT-PCR from D3 to D5 with immunocytochemistry at D5. We also inhibited BMP activity using 20 ng/mL NOGGIN at the same time-intervals (**Figure 4A**). On D3, early BMP4 exposure significantly increased *FOXF1* and *HAND1*, with NOGGIN causing the opposite effect, supporting an early role of BMP signaling in LPM fate decision^22^. On D4, both early and mid BMP4 exposures markedly increased *FOXF1* and *HAND1*, while *MEOX1* was downregulated. The inverse was true after NOGGIN treatments, joined by increased PAX3 expression. In contrast, *OSR1* was downregulated in both BMP4 and NOGGIN treatments (**Figure 4B**) and might be due to its shared expression in LPM and IM progenitors early on during development, after which *Osr1* becomes restricted to the IM lineage at later stages^3,35^. On D5, BMP4 or NOGGIN addition showed clear opposing effects on *FOXF1*/*HAND1* and *PAX3*/*MEOX1* expression as seen on D3 and D4, with the strongest effect generated in early and mid-timed pulses. The more restricted IM markers, *PAX8* and *PAX2* became expressed by D5, as well as later IM markers *WT1, SIX2* and *LHX1*. Interestingly at D5, early and mid BMP4 addition significantly reduced *PAX8* and *PAX2*, but late BMP4 addition did not. Moreover, late BMP4 stimulation was permissive to keeping the IM lineage intact, and significantly increased *OSR1* and *LHX1* expression with a concomitant decrease in PXM markers. Importantly, IM markers *OSR1, PAX2* and *PAX8* were reduced across all NOGGIN treatment windows, including at D5 and joined by *WT1* and *SIX2* reduction, indicating that BMP signaling is required for both early LPM and later IM specification (**Figure 4B**).

**Figure 4.**
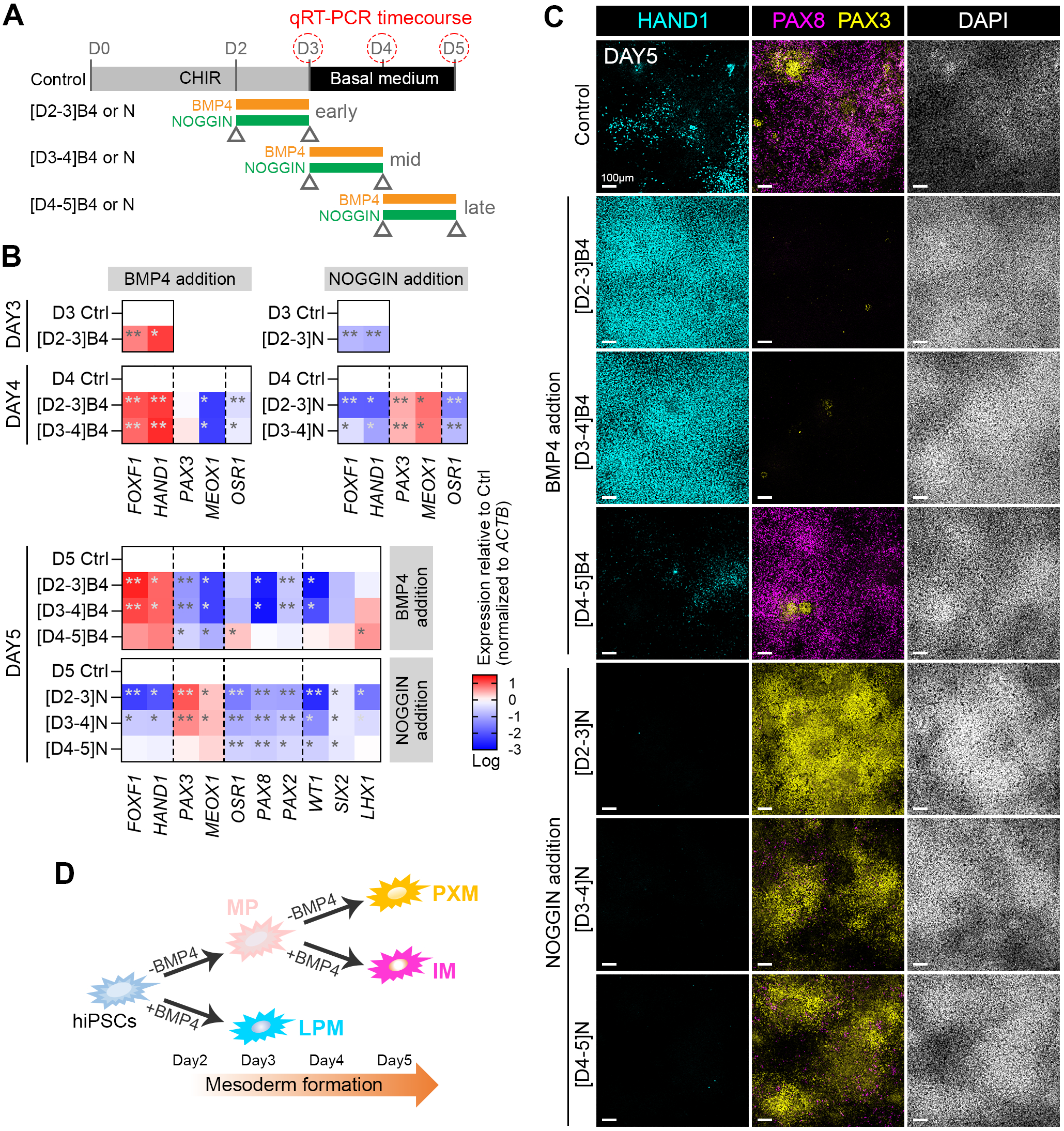
BMP4 exerts temporally distinct effects on LPM and IM specification. **(A)** Schematic showing experiment setup: hiPSCs were treated with 4 μM CHIR99021 (CHIR) for 72 h, followed by a 24 h treatment of either 5 ng/mL BMP4 (B4, orange) or 20 ng/mL NOGGIN (N, green) on D2 ([D2-3], early), D3 ([D3-4], mid), or D4 ([D4-5], late). **(B)** qRT-PCR performed 24 h after factor addition for PXM, IM, and LPM markers in induced cells after BMP4 or NOGGIN treatments. Heatmaps show Log expression versus D3, D4 or D5 control cells. Expression is the average from minimum five independent experiment, with statistical significance shown by asterisks (*, p<0.05; **, p<0.001 in a standard t-test, condition versus control). **(C)** Immunofluorescent staining at D5 for PAX3, PAX8, and HAND1 proteins in different experimental conditions. Nuclear counterstain with DAPI. **(D)** Schematic summary of BMP4 signal modulation and observed mesodermal lineage patterns.

Consistent with these transcriptional changes, immunofluorescent staining showed predominantly HAND1^+^ LPM cells were generated by both early and mid BMP4 stimuli. In contrast, late BMP4 treatment reduced HAND1^+^ cells. Instead PAX8^+^ IM cells had maintained or even slightly expanded in culture compared to the mixed mesoderm of the control (**Figure 4C**). On the other hand, NOGGIN markedly inhibited HAND1^+^ LPM cells and PAX3^+^ cells dominated the cultures (**Figure 4C**), reflecting the expansion of the PXM lineage under BMP inhibition^22^. Since D5 control cultures express *PAX8* / PAX8 (**Figure 1B** and **4C**), a late BMP4 stimulus might have maintained or slightly expanded this IM population, judged by D5 *OSR1* expression (**Figure 4B**), potentially from the mixed mesoderm seen in D4 sub-clustering (**Figure 3G-I**). Moreover, under late NOGGIN exposure, more PAX8^+^ cells had formed compared to early and mid conditions, suggesting that BMP signaling at D4 positively enhanced IM formation, likely from cells that cluster together with D4 PXM progenitors. Taken together, these results demonstrate a temporal switch in BMP4 responsiveness, reflecting the temporal *BMP4* expression pattern in our scRNA-seq dataset, whereby early BMP4 signaling drives LPM specification and late BMP4 exposure preferentially promotes IM differentiation from a mixed mesoderm progenitor population (**Figure 2E** and **4D**).

### Early BMP4 stabilizes LPM fate and late BMP4 directs mesoderm plasticity

To further test this temporal role of BMP signaling, we examined the response of mesoderm progenitors in 5 ng/mL BMP4 followed by its later inhibition with 20 ng/mL NOGGIN. Since the level of BMP4 has been shown to drive mediolateral fate decisions *in vivo*^37,54^, we also performed the opposite experiment in which we tested a range of BMP4 concentrations after initial BMP signal inhibition to investigate the role of *BMP4* in the fate decision step of the later mixed mesoderm progenitor population (**Figure 5A-B** and **S8A**). Early BMP4 treatment resulted in increased *FOXF1* and *HAND1* expression and reduced *PAX3, MEOX1, OSR1* and *PAX8* expression, confirming early additional BMP4 stabilizes LPM fate at the expense of PXM/IM formation on D3 (**Figure 5C**). Immunofluorescent staining further showed most cells had adopted a HAND1^+^ LPM-like identity by D3 and that subsequent BMP inhibition did not substantially alter this expression pattern (**Figure 5D-E**). This indicates that early BMP exposure pivots as well as stabilizes cells in a stable LPM state.

**Figure 5.**
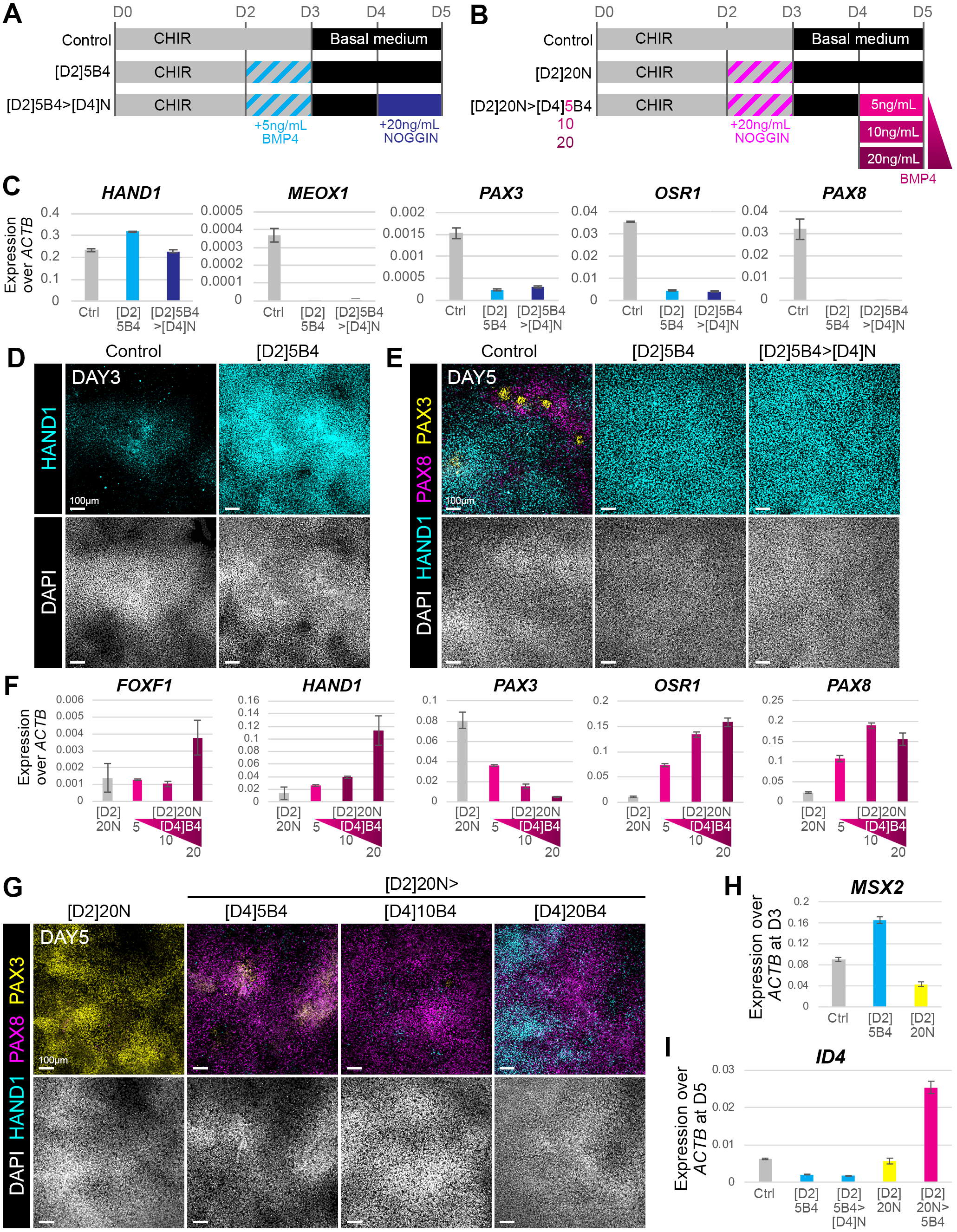
Sequential BMP modulation affects mesodermal lineage specification. **(A-B)** Schematics showing experiment setups in which hiPSCs were treated with 4 μM CHIR99021 (CHIR) for 72 h, followed by basal medium culture from D3 (Control condition). **(A)** An additional 24 h [D2-3] treatment of 5 ng/mL BMP4 (B4) was either followed by basal medium or a 24 h [D4-5] 20 ng/mL NOGGIN (N) treatment. **(B)** Reverse to experimental setup in A, an initial 24 h [D2-3] treatment of 20 ng/mL NOGGIN was followed by basal medium or a 24 h [D4-5] BMP4 pulse. A range in BMP4 concentrations was tested (5, 10, or 20 ng/mL). **(C)** qRT-PCR analysis on D5 for *PAX3* and *MEOX1* (PXM); *OSR1* and *PAX8* (IM) and *FOXF1* and *HAND1* (LPM) markers in experiment shown in A (one representative experiment shown out of four independent experiments with similar outcome). **(D-E)** Immunofluorescent staining for PAX3, PAX8, and HAND1 proteins in experiment A, on D3 **(D)** and D5 **(E). (F)** qRT-PCR analysis on D5 for *PAX3* and *MEOX1* (PXM); *OSR1* and *PAX8* (IM) and *FOXF1* and *HAND1* (LPM) markers under the indicated treatment conditions in experiment shown in B (one representative experiment shown out of two independent experiments with similar outcome). **(G)** Immunofluorescent staining of cells under the indicated treatment conditions in experiment B against PAX3, PAX8 and HAND1 on D5. Nuclear counterstain with DAPI. **(H)** Expression of *MSX2* at D3 in different treatment conditions, shown in A and B. **(I)** Expression of *ID4* at D5 in different treatment conditions, shown in A and B. Column color denotes the predominant fate (grey, control; cyan, LPM; yellow, PXM; pink, IM). Data is the average expression of two independent experiments.

In the reverse experiment, mesodermal cells - initially directed away from BMP-driven LPM differentiation by early NOGGIN addition - had retained the capacity to respond to later BMP4 stimulation. D2 NOGGIN addition efficiently suppressed LPM markers *FOXF1* and *HAND1* and augmented *PAX3* and *MEOX1* expression by D5 (**Figure 5F** and **S8B-C**). Interestingly, on D4 this cell population still expressed MP markers such as TBX6 and CDX2 (**Figure S9**), consistent with a state of retained developmental plasticity. Remarkably, reintroduction of BMP4 at D4 redirected these mixed mesoderm progenitors in a dose-dependent manner. Moderate BMP4 concentrations (5-10 ng/mL) reduced PXM-associated *PAX3* expression, upregulated IM markers *OSR1* and *PAX8* and subsequently induced the nephrogenic marker PAX2 at D9 and D12 (**Figure S8D**). In contrast, 20 ng/mL BMP4 increased LPM marker expression, alongside *OSR1*^+^ *PAX8*^+^ IM cells at D5 (**Figure 5F**), however, higher concentrations of BMP4 reduced *OSR1*^+^ *PAX2*^+^ *PAX8*^+^ IM specification (**Figure S8D**). D2 NOGGIN-treated cells did not contain any obvious HAND1^+^ population on either D3 or D4, suggesting that later, higher levels of BMP4 from D4 to D5 can convert TBX6^+^ CDX2^+^ mesoderm progenitors to LPM-like cells (**Figure S9**). Additional immunofluorescent staining on D5 showed NOGGIN inhibition induced relatively pure PAX3^+^ PXM-like cells, whereas 5 ng/mL BMP4 increased the amount of PAX8^+^ cells together with residual PAX3^+^ populations, indicating a mixed transition from the mixed mesoderm progenitor pool. At 10 ng/mL BMP4, IM specification was strongly favored with most cells expressing PAX8 along with small residual populations of PAX3^+^ PXM-like and HAND1^+^ LPM-like cells. In contrast, 20 ng/mL BMP4, increased HAND1^+^ at the expense of PAX8^+^ cells (**Figure 5G**). Finally, to confirm our observations in scRNA-seq (**Figure 2F, 3D** and **3J**), we tested *MSX2* and *ID4* expression by qRT-PCR in the timed conditions that had largely biased to either LPM or IM fates (**Figure 5H-I**). We found upregulated *MSX2* in D3 LPM and increased *ID4* in D5 IM populations. Moreover, BMP4 at D4-5 increased *ID4* expression in a dose-dependent manner (**Figure S8E**).

Together, these results indicate that progenitors escaping early LPM specification retain the plasticity to form IM and LPM lineages in a BMP4 dose- and time-dependent manner. This further implies that the IM lineage is best induced within a defined time and dose-dependent window of BMP signaling.

### PXM progenitors exhibit time-dependent competence for BMP4-mediated repatterning

Given that transient BMP inhibition on D2 generated a PXM-like population competent for subsequent IM induction (**Figure 5F-G**), we next asked whether MPs under a stricter experimental condition that limited early LPM formation, could also be reprogrammed toward alternative mesodermal fates. To block early LPM and promote PXM specification^55–58^, hiPSCs were treated from D0-3 with 4 µM CHIR and 0.1 µM LDN193189, a BMP inhibitor, followed by 24 h pulse with 5 or 20 ng/mL of BMP4 at D3, D4 or D5. The day after BMP treatment, cultures were analyzed by qRT-PCR and immunofluorescent staining (**Figure 6A**). D3 BMP4 treatment still robustly induced HAND1^+^ LPM cells in both low and high BMP4 concentrations, indicating D3 MPs remain permissive for LPM specification (**Figure 6B-C**). In contrast, BMP inhibition without any BMP4 stimulus generated a predominant PAX3^+^ PXM population (**Figure 6D-E**). D4 BMP4 treatment, however, reduced both *PAX3* transcript and protein expression and concomitantly induced IM and LPM markers in a dose-dependent manner, with low BMP4 preferentially promoting PAX8^+^ IM cells and higher concentrations increasing co-induction of IM and LPM markers (**Figure 6B** and **6D**). In D5 BMP4 treatment, some PAX8^+^ cells were observed in the absence of detectable LPM markers (**Figure 6B** and **6E**), indicating PXM cells had lost competence for LPM specification while retaining the ability to generate IM.

**Figure 6.**
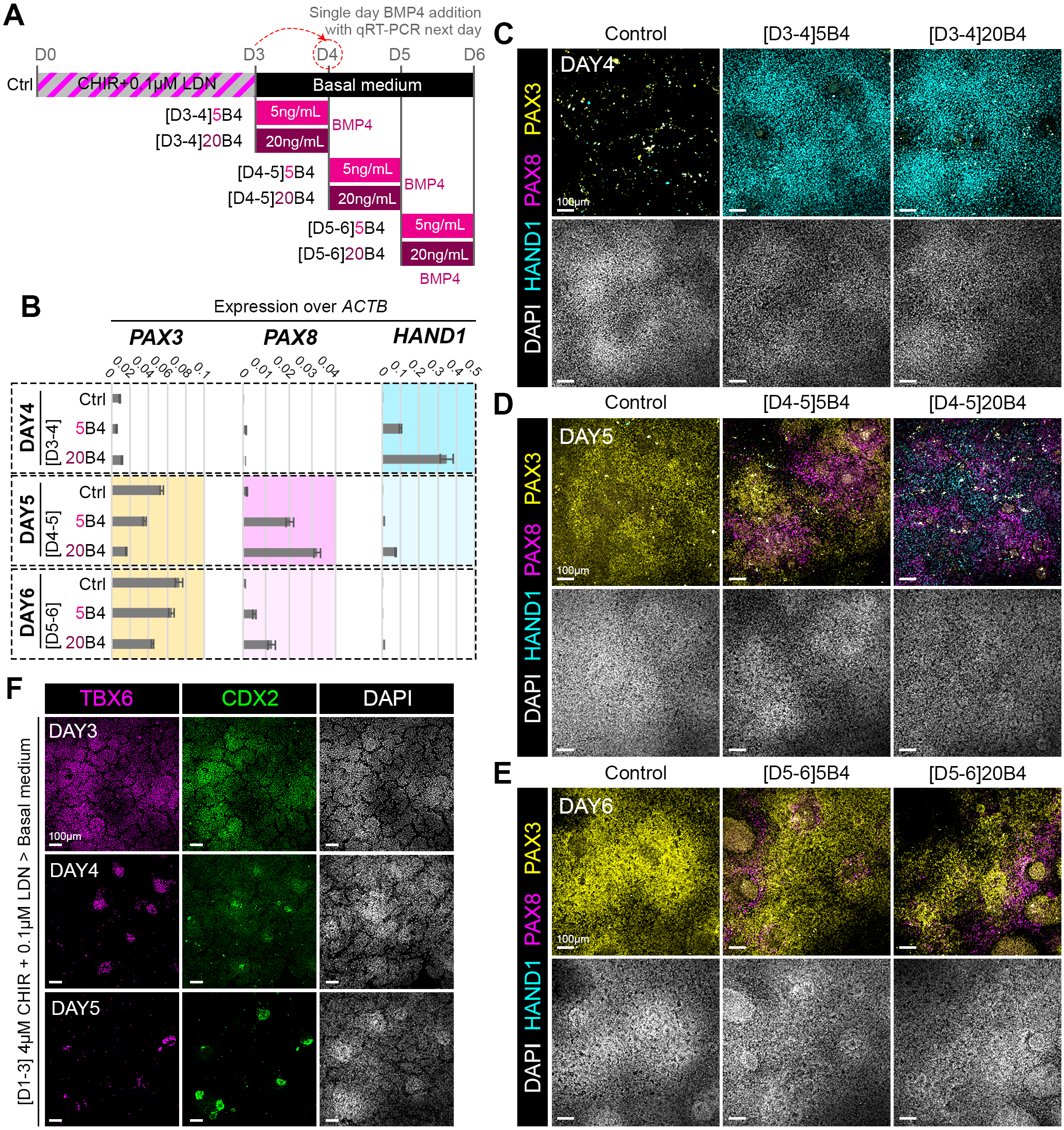
PXM progenitors exhibit time-dependent competence for BMP4-mediated repatterning. **(A)** Schematic showing experiment setup: hiPSCs were treated with 4 μM CHIR99021 (CHIR) and 0.1 μM LDN193189 (LDN) for 72 h, followed by basal medium culture from D3 (Ctrl condition). On D3, D4 or D5, cells were supplemented with a 24 h BMP4 pulse (5 or 20 ng/mL) and collected 24 h after BMP4 treatment. **(B)** qRT-PCR analysis on D4, D5 or D6 of *PAX3* (PXM), *PAX8* (IM) or *HAND1* (LPM). One representative experiment shown out of three independent experiments with similar outcome. **(C-E)** Immunofluorescent staining for PAX3, PAX8, and HAND1 proteins at D4 **(C)**, D5 **(D)** or D6 **(E)** under the indicated conditions. **(F)** Immunofluorescent staining for CDX2 and TBX6 proteins under control conditions at D3, D4 and D5. Nuclear counterstain with DAPI.

Notably, D4 BMP4 administration resulted in more robust IM induction, indicating an optimal temporal window for its specification. IM induction efficiency under this protocol, however, was lower than that following D2 BMP inhibition by NOGGIN (**Figure 5G**), likely reflecting both the reinforced PXM specification from prolonged LDN193189 treatment as well as the depletion of TBX6^+^ CDX2^+^ progenitors (**Figure 6F**). TBX6^+^ CDX2^+^ progenitors were abundant on D3, with TBX6 expression declining over time. In contrast, CDX2 expression persisted longer and formed CDX2^high^ core-like structures that progressively expanded and adopted posterior neural tube-like morphologies, characterized by the presence of SOX2 but absence of NCC marker SOX10. BMP4 treatment at D3, however, could maintain CDX2^low^ TBX6^+^ progenitors for longer (Figure S10). Interestingly, these CDX2^high^ structures further developed an ordered spatial organization with CDX2^high^ cells at their core, surrounded by TBX6^+^ cells and PAX8^+^ cells at the periphery, recapitulating the mediolateral organization of the posterior neural tube, presomitic mesoderm (PSM) and IM tissues observed during embryonic development (Figure S11).

Together, these results demonstrate a time-dependent shift in mesodermal competence in response to BMP4 signaling: early exposure favors LPM specification, intermediate exposure permits IM and LPM in a dose-dependent manner, and late exposure selectively supports IM differentiation from PXM-like cells.

### Temporal BMP modulation enables IM progression to nephrogenic tissue

Since D2 BMP inhibition by NOGGIN (20 ng/mL) followed by BMP4 re-activation on D4 (10 ng/mL) efficiently generated a substantial population of PAX8^+^ cells, we next sought to determine whether these cells represent functionally competent nephrogenic progenitors. To test this, D5 cells were further maintained in basal culture medium and grown until D12 to check their capacity for sustained kidney lineage progression and structural organization. We compared this condition to the control (three days in CHIR, which produces heterogeneous mesoderm; **Figure 1D**) and D2 NOGGIN-treated cells. In all conditions D12 cells had generated to some degree kidney-like structures expressing key nephric markers, including GATA3, LHX1, and WT1. Under our optimized protocol, however, organoids consistently exhibited more advanced maturation, characterized as the robust and orderly co-expression of GATA3, LHX1, and WT1, together with the emergence of segmented structural organization. Across four independent experiments, quantitative scoring revealed that 80% of organoids reached this well-patterned state under our temporal BMP modulation protocol. In contrast, the maturation was substantially lower in the control condition, with only 50% of organoids achieving comparable features following CHIR-only induced differentiation, and 33% under CHIR/NOGGIN-only treatment (**Figure 7A-B**). Together, these findings indicate that temporally controlled BMP signaling not only enhances the efficiency of IM-derived organoid formation but also promotes their structural maturation into organized kidney-like architectures.

**Figure 7.**
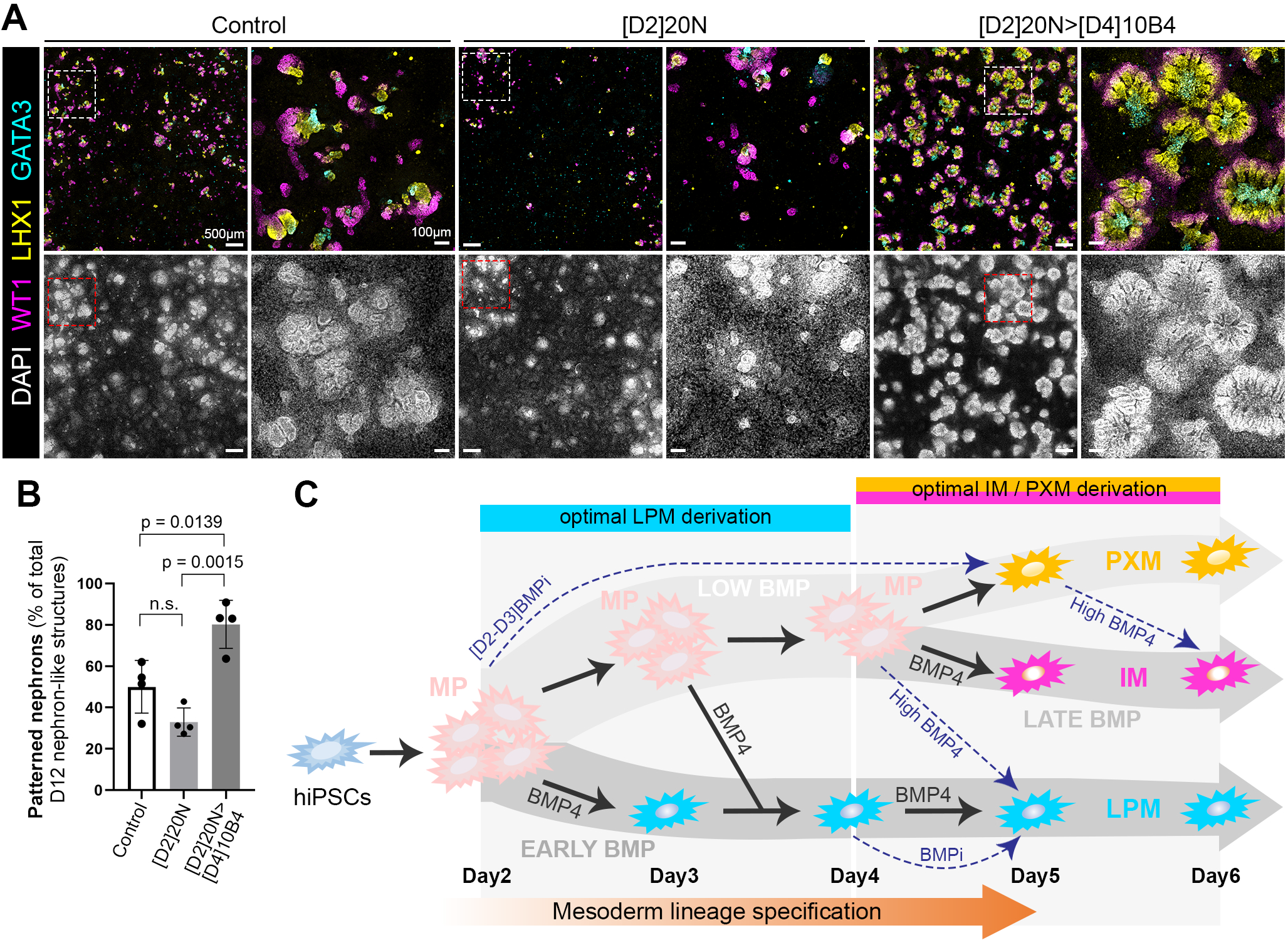
Temporal BMP modulation enables IM progression to patterned nephrogenic tissue. **(A)** Immunofluorescent staining against LHX1, GATA3, and WT1 proteins on D12. Whole-well (48-well plate, stitched orthogonal projection of Z-stack) imaging is shown on the left, with representative higher-magnification images on the right. Control, [D1-3] 4 µM CHIR followed by basal medium; [D2]20N, same as control but with an additional 24 h treatment of 20 ng/mL NOGGIN on D2; [D2]20N>[D4]10B4, same as [D2]20N condition with an additional 24h treatment of 10 ng/mL BMP4 on D4. **(B)** Quantification of patterned nephron organoid structures under the conditions in A, shown as a percentage of the total D12 nephron-like structures. Nephrons were characterized as mature/patterned if they presented correct alignment of WT1-LHX1-GATA3 expression. Columns represent the mean across four independent experiments (black dots) with error bars the standard deviation (p-values show statistical significance, calculated by a two-way ANOVA test; n.s., non-significant). **(C)** Model based on scRNA-seq data and experimental BMP modulation experiments during hiPSC differentiation to illustrate lineage relation, LPM-IM-PXM specification as well as the plasticity between them in response to timed BMP4 signaling. Black arrows show the postulated role of BMP4 driving both early LPM and late IM mesoderm lineage bifurcations. Blue dashed arrows represent mesoderm plasticity when BMP4 is added or inhibited by NOGGIN or LDN193189 (BMPi).

## DISCUSSION

Here, we presented a heterogeneous mesoderm differentiation system from hiPSCs that enables the coexistence of multiple mesodermal lineages, in which the fate distribution can be redirected by BMP modulation within a defined temporal framework. This work offers insight into early human mesoderm differentiation at the single cell level and provides a paradigm for the study of other dynamic signaling inputs that influence mesoderm patterning. Finally, by temporally modulating BMP signaling, we promoted IM specification and increased nephrogenic maturation.

### Temporal BMP4 signaling governs mesoderm lineage specification

Our scRNA-seq analyses show that under minimal CHIR-induced mesoderm induction, BMP4 is one of the earliest signals of mesodermal fate divergence at early days of differentiation (**Figure 2** and **3**). Studies in vertebrate embryos have demonstrated that graded BMP activity specifies distinct mesodermal fates, with high BMP promoting lateral identities and lower levels favoring more medial derivatives, including PXM and IM^37,38,54,59,60^. These findings support a quantitative model in which BMP signaling acts in a dose-dependent manner to control mesoderm patterning. While hiPSCs fate can be steered by increasing dose of BMP4 (**Figure 5, 6** and **S8**), confirming this dose-dependent framework, our findings also demonstrate that a BMP4 signal can be interpreted in a time-dependent manner during human iPSC-derived mesoderm formation. Here, we present a model in which early BMP4 exposure or inhibition promotes stable LPM or PXM fate, respectively (**Figure 4D** and **7C**), consistent with PXM-LPM divergence *in vitro* by BMP modulation^22,23^. We found a second BMP signal later diverges IM from PXM fate (Figure 4 and 5). As differentiation proceeds, mixed D3-4 PXM/IM mesoderm progenitors likely cluster together in pockets of low BMP signaling, only to be influenced by early LPM-derived BMP4 signaling later during culture. This temporal lineage progression mimics the consecutive tissues formed from the PS *in vivo*, as lateral fates emerge before medial fates in the early gastrula^7–9,11^. BMP responsiveness likely shifts as mesoderm progenitors mature and the ability to adopt LPM fate progressively declines and biases toward IM formation (**Figure 4**).

A potential mechanistic basis for this temporal shift is provided by our single-cell transcriptomic analysis. At D3, the extensive overlap between *BMP4*, its downstream target *MSX2* with canonical LPM markers *FOXF1* and *HAND1* suggests that BMP signaling is actively engaged during the establishment of LPM fate^22,53^. By D4, although *BMP4* expression is reduced - potentially reflecting changes in WNT signaling following CHIR withdrawal - *MSX2* expression remains strongly associated with LPM markers, indicating that the downstream transcriptional responses to BMP signaling persist despite reduced ligand availability (**Figure 3**). At the same time, *ID4* becomes expressed in a subset of cells co-expressing IM markers such as *OSR1* and *PAX8*, suggesting an alternative BMP-responsive transcriptional program is associated with IM specification. From D5 onwards, *BMP4* was broadly distributed across both LPM and IM populations, however, *MSX2* and *ID4* remained segregated along LPM and IM clusters, respectively. Although we did not directly test the functional roles of these downstream effectors during mesoderm lineage specification, it suggests that lineage-specific interpretation of BMP signaling is not determined by ligand distribution per se, but rather by distinct downstream transcriptional programs engaged over time.

### Plasticity of mesoderm progenitors enables lineage repatterning

By challenging our *in vitro* differentiation through timed BMP modulation, we revealed a temporally restricted window of mesodermal plasticity. Whereas the D4 LPM population could not be reverted to more medial fates (**Figure 5C-E**), suggesting lateral fate commitment had occurred by that point. In contrast, mesoderm subjected to transient BMP inhibition before D3 acquired a permissive, plastic state, enabling subsequent redirection toward LPM fates upon BMP re-exposure (**Figure 5F-G** and **6**). This plastic cell state remained in culture but decreased over time as D4 cells remained responsive to BMP signaling in a dose-dependent manner to convert to either IM or LPM fate (**Figure S8**). At D5, these cultures were largely defined by a PAX3^+^ PXM identity with few TBX6^+^CDX2^+^ progenitors left (**Figure 6F** and **S9**). Under low BMP signaling, such TBX6^+^CDX2^+^ progenitors, reflecting the PSM *in vivo*^61^, are likely largely predisposed toward a PXM trajectory and would upregulate PAX3 upon continued differentiation in basal culture conditions, corroborated by our experiments in LDN193189 or NOGGIN without further BMP treatment (**Figure 4-6**). Therefore, we assume that these PAX3^-^ cells represent not fully PXM-committed progenitors and remain amenable to repatterning. Such plasticity has been confirmed *in vivo* through various transplantation studies in mouse and avian embryos in which mesoderm fate can be re-directed by the local environmental cues of their new environment^12,18,48,62–64^, and attributed at least in part due to BMP signaling^18,38,48^. Signals derived from the PXM are required for *Pax2* induction and IM formation, and disruption of the interface between PXM and IM domains impairs nephric specification^64^. The different penetrance in single *Tbx6* and *Wnt3a* versus *Tbx6:Wnt3a* compound mutant embryos^65^, or the opposing effect seen in *Foxc1:Foxc2* compound mutant mouse compared to *Foxc1/Foxc2* overexpression in chick embryos^19^, further support that PXM and IM fates as well as their genetic regulation are closely linked, at least when the anterior somites, pro- and mesonephros form during early gastrulation. We show that during hiPSC differentiation, IM progenitors form in close relation to PXM progenitors but are already distinct at the single cell level (**Figure 2 and 3**). Within this context, BMP signaling continues to act in a dose-dependent manner with moderate levels favoring IM differentiation and higher levels retaining partial LPM-inducing capacity.

Our *in vitro* differentiation system reproducibly generates three mesoderm lineages and IM cells were often positioned between PXM and LPM populations, reflected by the spatial distribution of PAX2/8 between PAX3 and HAND1 expression (**Figure 1 and 5**). PAX8^+^ cells were also observed adjacent to CDX2^+^SOX2^+^ core-like structures surrounded by TBX6^+^ progenitors, partially recapitulating the medial-to-lateral organization of embryonic mesoderm (**Figure S10 and S11**). These findings suggest that IM specification is not solely determined by BMP4 dosage and timing but also depends on continued local interactions between these neighboring mesodermal populations. Taken together, our findings support a model of progressive lineage restriction, in which mesodermal competence is gradually constrained over developmental time through both loss of multipotent progenitors and reinforcement of lineage-specific transcriptional programs, thereby defining stage-dependent responses to BMP4 signaling (**Figure 7C**).

### BMP4 efficiently drives functional IM specification but requires additional signaling to enhance nephrogenic output

BMP4 is a well-established regulator of IM^37^ and kidney development^66^. *In vivo* studies have shown that BMP signaling from adjacent tissues is required for nephric duct formation and regulation of renal progenitor gene expression, supporting roles for BMP activity in both early IM specification and subsequent nephrogenic development^67^. Temporal modulation of BMP signaling efficiently induced early *OSR1*^+^ IM identity and well-patterned organoid structures by D12 under basal conditions (**Figure 5G** and **7A-B**). However, *PAX2* expression did not substantially increase at the population level (**Figure S8B**), suggesting that BMP4 promotes patterning but is insufficient to sustain further nephrogenic progenitor expansion.

One key limitation to achieve fully selective IM specification *in vitro* is likely due to the initial mesodermal heterogeneity we set out to achieve. Our intentional strategy employs only moderate WNT activation to reflect the mid-PS identity^10^ and generates diverse mesoderm lineages to enable for the analysis of lineage diversification and signaling-dependent fate decisions. In contrast, previously reported kidney organoid protocols typically achieve more IM tissues, using higher or longer WNT stimulation^26^. These protocols might instead induce more posterior PS or later-stage posterior identities to provide a shortcut to more homogeneous, later stage IM and kidney cell specification. This is supported by lineage tracing in mouse, showing the metanephric mesenchyme is at least in part derived from later stage *Tbx6*^+^ *Sox2*^+^ neuromesodermal-like progenitors^35^. Taken together, although sequential BMP modulation promoted IM marker expression and supported organoid formation in this work, complete lineage purification was not achieved, with persistent PXM and LPM populations observed, even by D12 (**Figure S8**). This indicates that early patterning of the PS constrained downstream lineage specificity. Thus, while our system is well suited for dissecting temporal and signaling mechanisms of mesoderm specification, it is less optimal for generating purified nephrogenic populations. Instead, it can be utilized to refine early patterning conditions, including modulation of WNT or FGF dosage or integration of additional cues, to improve IM enrichment and lineage specificity. In this regard, our scRNA-seq data revealed enrichment of FGF family members *FGF8* and *FGF17* as well as the WNT inhibitor *DKK1* during early mesoderm specification (**Figure 3F**). Also, the more established roles FGF signaling, including *Fgf9*^68^, or combination with other BMPs such as *Bmp7*^69,70^ (Figure S5A) could be reexamined in our system to robustly reinforce the nephrogenic program. Such targeted, sequential signaling adjustments will not only resolve remaining questions of human mesodermal competence but also improve the quality and reproducibility of hPSC-derived kidney organoids.

## AKNOWLEDGEMENTS

We thank Chie Fukui, Chiyomi Ito and Maiko Takahashi for technical support. We thank Anestis Tsakiridis, Anahí Binagui-Casas, Rie Ajima and Val Wilson for their critical review of the manuscript. We thank all members of the Laboratory for Human Organogenesis for their valuable feedback and discussions. This work was supported by JSPS KAKENHI Grant-in-Aid for Young Scientists (A) Grant Number JP17H05006.

## AUTHOR CONTRIBUTIONS

Conceptualization: W.Z., M.T.; Investigation: W.Z.; Validation: W.Z.; Formal Analysis: W.Z., F.J.W.; Visualization: W.Z., F.J.W.; Writing – Original Draft: Z.W., F.J.W.; Writing – Review & Editing: Z.W., F.J.W., M.T.; Supervision: M.T.; Funding Acquisition: M.T.

## METHODS

### Resource Availability

#### Lead Contact

Further information and requests for resources should be directed to Minoru Takasato.

#### Materials Availability

This study did not generate new unique reagents.

## Data and Code Availability

Single-cell RNA sequencing data generated in this study will be deposited in GEO (Gene Expression Omnibus) and will be available upon publication.

## Experimental Model and Subject Details

### hiPSC Culture

The human induced pluripotent stem cell (hiPSC) line CRL1502 clone C32 (provided by E.J. Wolvetang, The University of Queensland, Australia) was used in this study. Cell line authentication was performed prior to use. hiPSCs were maintained in StemFit® AK02N/Basic03 medium (Ajinomoto; Cat# AK02N), supplemented with 1X Antibiotic-Antimycotic (Thermo Fisher Scientific; Cat# 15240062) under standard culture conditions. Cells were dissociated using TrypLE™ Select Enzyme (Thermo Fisher Scientific; Cat# 12563011) prior to seeding for all differentiation experiments.

## Method Details

### Mesoderm Induction

Monolayer, feeder-free differentiation was performed using STEMdiff™ APEL™ 2 medium (STEMCELL Technologies; Cat# ST-05275) supplemented with 3% Protein-free Hybridoma Medium (PFHM-II; Thermo Fisher Scientific; Cat# 12040077) and 1X Antibiotic-Antimycotic. hiPSCs were seeded onto iMatrix-511-coated culture plates and cultured overnight in StemFit medium supplemented with 10 μM ROCK inhibitor Y-27632. For single-cell RNA sequencing experiments, cells were seeded at a density of 3.5 × 10^4^ cells/cm^2^. Mesoderm induction was performed by treating hiPSCs with CHIR99021 (R&D Systems; Cat# 4423) for 72 h. The concentration was optimized (**Figure S1**) with 4 μM CHIR99021 used in all experiments. BMP signaling inhibitors including NOGGIN (R&D Systems; Cat# 6057-NG) and LDN193189 (TOCRIS; Cat# 6053) were applied as indicated.

### Single-Cell RNA Sequencing

Mesodermal cells derived from hiPSCs were dissociated into single cells using Bacillus licheniformis protease (Sigma-Aldrich; Cat# P4860) for 10 min on ice. Approximately 3,000 cells per sample were collected at six time points (D3-D7 and on D9 of differentiation). After quality control to remove dead cells, we obtained 13,967 cells in total. Single-cell capture, RNA purification, and reverse transcription were performed using the Chromium™ Single Cell Controller and Chromium Single Cell 311 Reagent Kits v3 (10x Genomics) following the manufacturer’s instructions. Sequencing libraries were prepared according to the manufacturer’s protocol and sequenced on an Illumina NovaSeq 6000 platform (GENEWIZ). A total of 3.13 × 10^9^ sequencing reads were generated across six samples, corresponding to an average sequencing depth of approximately 1.7 × 10^5^ reads per cell.

### Single-Cell RNA-Seq Data Analysis

Raw sequencing data were processed using the Cell Ranger pipeline (10x Genomics) with default parameters to generate gene expression matrices. Downstream analyses were performed using Seurat (v3.2.3)^71^. Cells were filtered based on quality control metrics. Cells with fewer than 2,500 detected genes or more than 8,000 detected genes, as well as those with greater than 15% mitochondrial gene expression, were excluded from downstream analyses. Gene expression values were normalized using the “LogNormalize” method with a scaling factor of 10,000. Highly variable genes (n = 2,000) were identified using the “FindVariableFeatures” function with the “vst” method. To reduce technical variation, gene expression data were scaled using the “ScaleData” function while regressing out the effects of mitochondrial gene expression, total UMI counts (nCount_RNA), and cell cycle scores (S phase and G2/M phase genes). Principal component analysis (PCA) was performed using the highly variable genes. The number of principal components (PCs) retained for downstream analysis was determined based on the elbow plot, by inspecting the point at which the standard deviation of PCs began to plateau. The top 16 PCs were selected based on the elbow plot and inspection of the variance distribution and were subsequently used for clustering and dimensionality reduction.

Cells were clustered using the “FindNeighbors” and “FindClusters” functions with a resolution parameter of 0.5. Uniform Manifold Approximation and Projection (UMAP) was performed using the same principal components (dims = 1:16) for visualization. Cluster-specific marker genes were identified using the “FindAllMarkers” function with the Wilcoxon rank-sum test, considering only genes that were expressed in at least 25% of cells in a cluster and with a log2 fold change greater than 0.25. Gene expression patterns across clusters were visualized using “DoHeatmap” and “FeaturePlot” functions. All analyses were performed using R (v4.1.3) and Seurat (v3.2.3)^71,72^.

### Immunofluorescence Staining

Cells cultured in 48-well plates were fixed with 4% paraformaldehyde for 5 min at room temperature and washed three times with DPBS. Samples were incubated with primary antibodies overnight at 4⍰°C, followed by incubation with fluorophore-conjugated secondary antibodies for 1 h at room temperature, with DPBS washes between. Fluorescence images were acquired using a laser scanning confocal microscope with GaAsP detectors (LSM 800; Zeiss). When entire wells were imaged, we used the “stitching” algorithm in ZEN software (Zeiss). All immunofluorescent staining images are orthogonal projections of Z-stacks of representative areas of the cultures. Details of primary antibodies are provided in Table S1.

### Quantitative RT-PCR

Total RNA was extracted using the NucleoSpin® RNA kit (TaKaRa Bio; Cat# 740955.250). cDNA synthesis was performed using PrimeScript™ RT Master Mix (TaKaRa Bio; Cat# RR036A). Quantitative reverse transcription polymerase chain reaction (qRT-PCR) was carried out using TB Green™ Premix Ex Taq™ II (TaKaRa Bio; Cat# RR820A) on a QuantStudio™ 5 Real-Time PCR System. Primer sequences are listed in Table S2.

### Quantification and Statistical Analysis

Statistical analyses were performed using R (v4.1.3). Differential gene expression analysis for single-cell RNA sequencing data was conducted using the Seurat package with the Wilcoxon rank-sum test. P values were adjusted for multiple testing using the Bonferroni correction. Genes with an adjusted p value < 0.05, expression in at least 25% of cells, and a log2 fold change > 0.25 were considered differentially expressed. For nephron organoid quantification, images of four independent experiments were manually counted in Photoshop (Adobe). Only structures that contained proper proximal to distal expression of WT1-LHX1-GATA3 proteins were scored as properly patterned. Graph and statistics (two-way ANOVA) were done in Prism9 (v9.5.1, Graphpad).

