## Supplemental Figures for "Time-dependent BMP4 signaling directs lineage specification in human mesoderm"

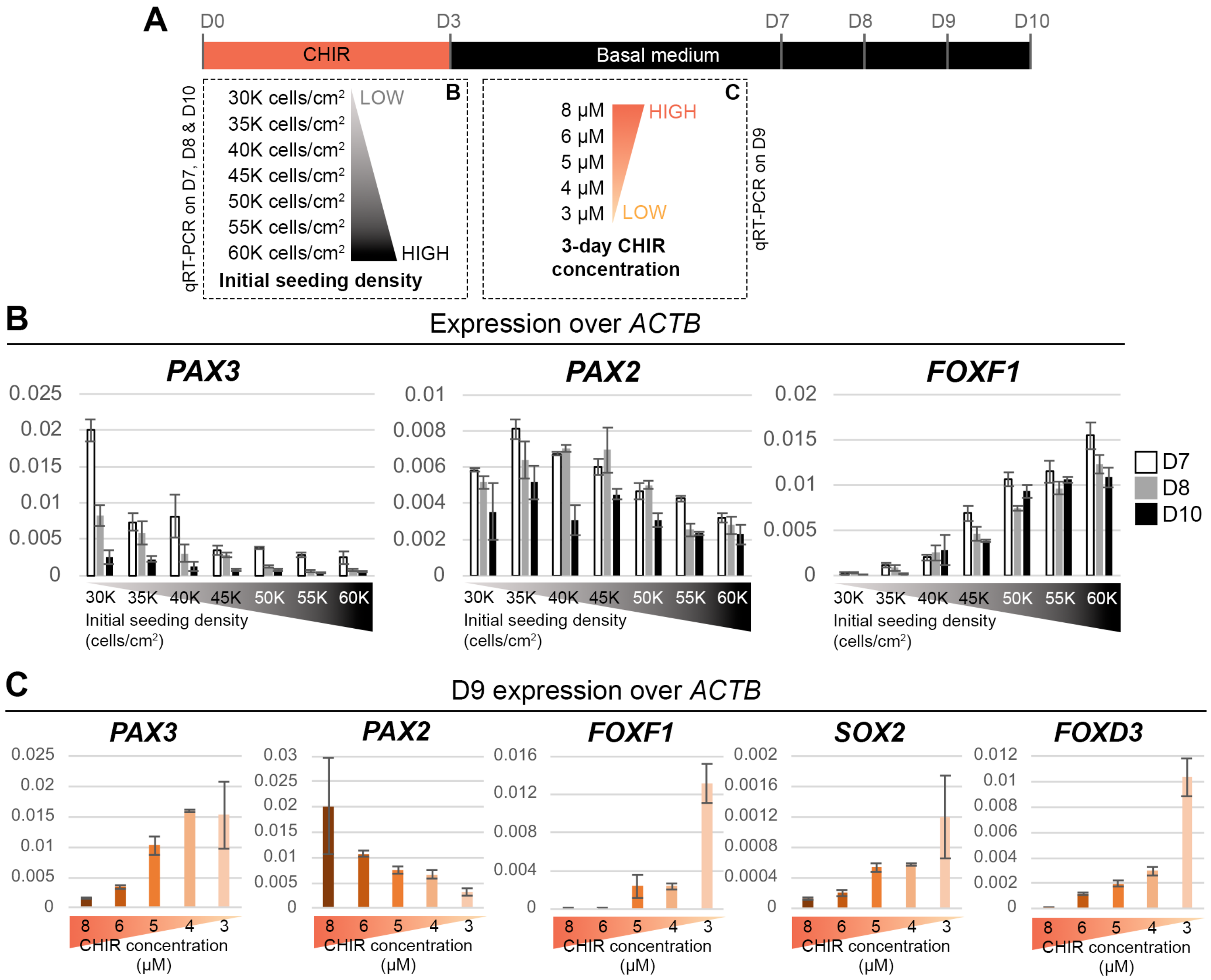


**Figure S1. Cell seeding density and CHIR99021 concentration affect mesoderm lineage proportions.** (**A**) Experimental design for cell seeding density and CHIR99021 concentration optimization during hiPSC differentiation. In one experiment, cells were seeded at densities ranging from 30K to 60K cells/cm² (K=1000). In a separate experiment, different CHIR99021 concentrations (8, 6, 5, 4 and 3 µM CHIR) were applied during the first three days of differentiation, followed by culture in basal medium from D4 for an additional six days. (**B-C**) Gene expression by qRT-PCR analysis tested for adequate PXM, IM and LPM outcomes. (**B**) Expression of *PAX3* (PXM), *PAX2* (IM), and *FOXF1* (LPM) under the indicated cell seeding density conditions at D7, D8 and D10. (**C**) Expression on D9 of *PAX3* (PXM), *PAX2* (IM), and *FOXF1* (LPM) and *SOX2* (neural) and *FOXD3* (neural crest) under the indicated CHIR concentrations.

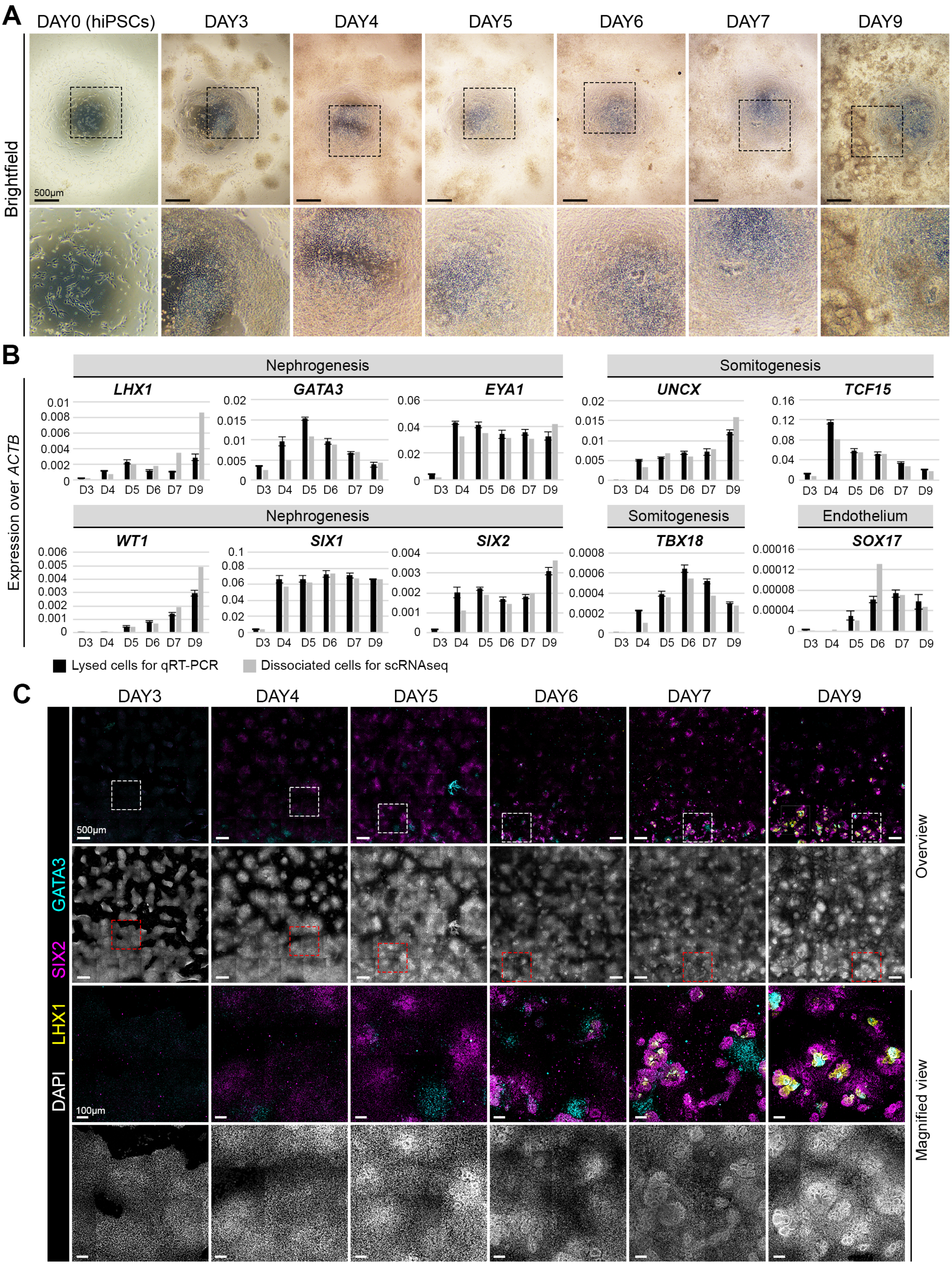


**Figure S2. hiPSC differentiation generates heterogeneous mesoderm for scRNA-seq analysis.**

**Figure S2. hiPSC differentiation generates heterogeneous mesoderm for scRNA-seq analysis.**
(**A**) Bright-field images of differentiating hiPSC cultures on D0, D3-D7 and on D9, used for scRNA-seq. (**B**) qRT-PCR analysis of markers associated with nephrogenesis, somitogenesis and endothelium in parallelly-grown differentiation showing dissociated cells, used for qRT-PCR validation (grey) and in collected samples used for scRNA-seq (black). (**C**) Immunofluorescent staining of nephrogenic markers LHX1, SIX2 and GATA3 on D3-D7 and on D9 of hiPSC differentiation. Images are stitched orthogonal projections of confocal stacks of one well (48-well plate).

A

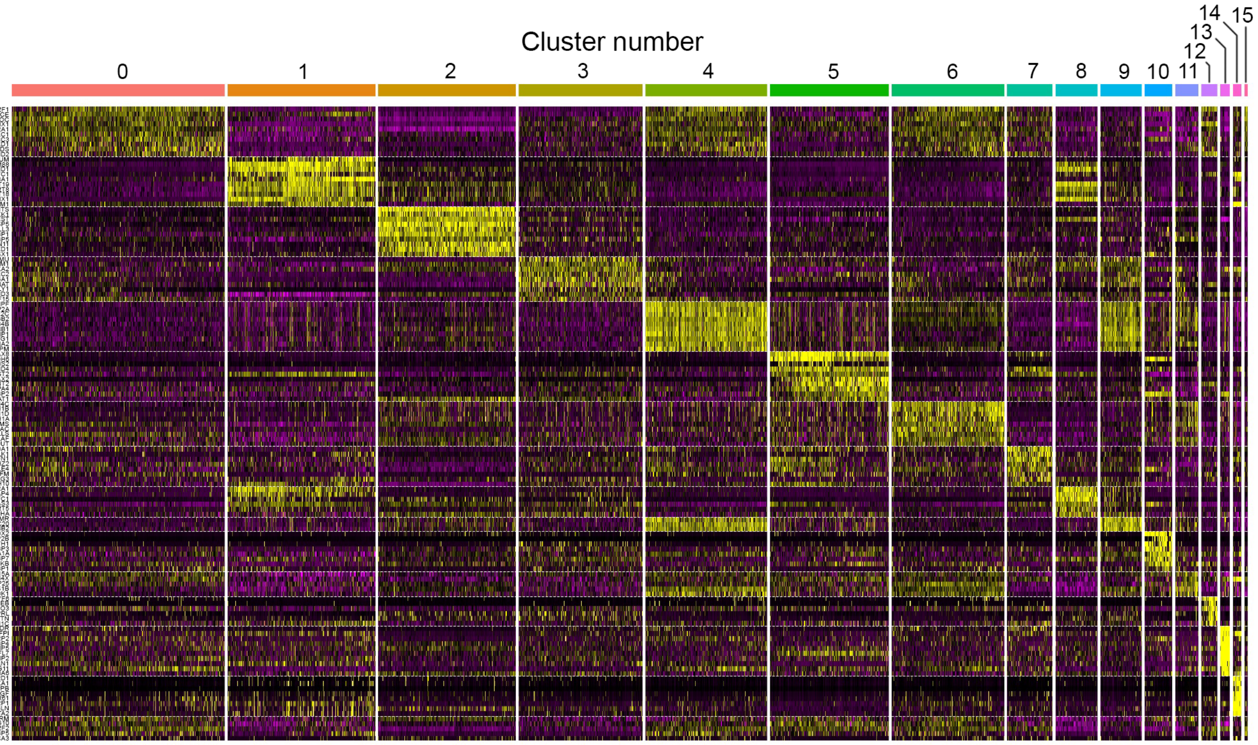


B

**TOP 10 DEGs in unsupervised hierarchical clustering**

| **0** | **1** | **2** | **3** | **4** | **5** | **6** | **7** |
|---|---|---|---|---|---|---|---|
| *NR2F1* | *LUM* | *NTS* | *NMU* | *CENPF* | ***PAX8*** | *HIST1H4C* | *PHLDA1* |
| *PCOLCE* | *TMEM88* | *DKK1* | *HOTAIRM1* | *TOP2A* | *CDH6* | *HIST1H1B* | *DLK1* |
| *BOC* | ***HAND1*** | *MEST* | *ANXA2* | *UBE2C* | *G0S2* | *HIST1H1D* | *FN1* |
| *SIX1* | *ACTC1* | *SP5* | ***FOXC2*** | *HMGB2* | ***ID4*** | *HIST1H1A* | *TSHZ2* |
| *COL2A1* | *COL3A1* | *DLL3* | *COL14A1* | *TUBB4B* | *BST2* | *TYMS* | *TLE4* |
| ***FOXC1*** | *KRT19* | *RBP1* | *NNAT* | *CCNB1* | ***PAX2*** | *HIST2H2AC* | *NEFM* |
| *IGDCC3* | *KRT8* | *FABP5* | ***RIPPLY1*** | *ARL6IP1* | *SPINT2* | *HELLS* | *MEG3* |
| *FOXD1* | *KRT18* | *CLDN11* | *ID3* | *PTTG1* | *DPPA4* | *PCLAF* | *RDH10* |
| *GMDS* | *LIX1* | *NKD1* | ***TCF15*** | *KPNA2* | *IGFBP2* | *DUT* | |
| *ID2* | *TPM1* | *MSX1* | | *ASPM* | *SAT1* | | |

| **8** | **9** | **10** | **11** | **12** | **13** | **14** | **15** |
|---|---|---|---|---|---|---|---|
| *HTRA1* | *HMMR* | ***SOX2*** | *TMSB15A* | *MYF6* | ***KDR*** | *ANKRD1* | *SRM* |
| *MFAP4* | *CDC20* | ***TFAP2B*** | *TMSB4X* | *NEB* | *TFPI* | *ANXA1* | *PEG10* |
| *CFC1* | *CCNB2* | *TTYH1* | *SNRNP25* | *ENO3* | *NRP2* | *NPPB* | *NR2F2* |
| *HOXB-AS3* | | *FGFBP3* | *CKS1B* | *PRL* | *IGFBP4* | *CTGF* | *IGFBP5* |
| *ART5* | | *TUBA1A* | *CDK1* | *TTN* | *VAMP5* | *CYR61* | *CA3* |
| *LDHA* | | *FABP7* | | *CDKN1C* | *EGFL7* | *CSRP1* | |
| | | *CKB* | | | *RAMP2* | *TAGLN* | |
| | | *CRABP1* | | | *HAPLN1* | *ACTA2* | |
| | | | | | *GNG11* | | |
| | | | | | *S100A6* | | |

**Figure S3. Characterization of cell clusters in unsupervised hierarchical clustering.**
(**A-B**) Heatmap (**A**) and summary table (**B**) of the top ten marker genes for each cluster (*cluster 0-15*). Markers indicative of the PXM, IM, LPM, endothelium or NCC are shown in bold.

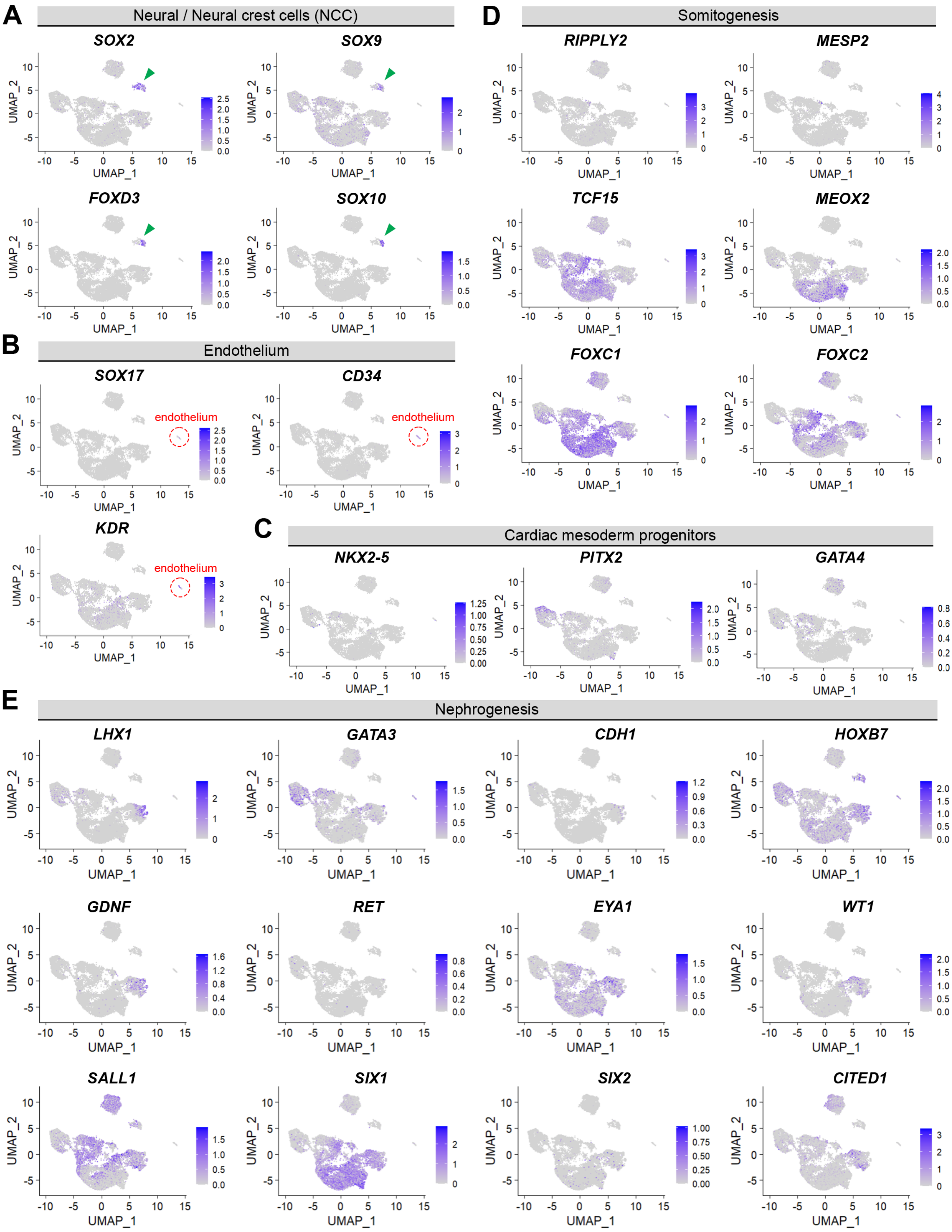


**Figure S4. UMAP feature plots in D3-9 cells.**
(**A-D**) UMAP feature plots during D3-9 hiPSC differentiation showing single genes indicative of neural / neural crest (NCC; **A**), endothelium (**B**), cardiac mesoderm (**C**), somitogenesis (**D**) or nephrogenesis (**E**).

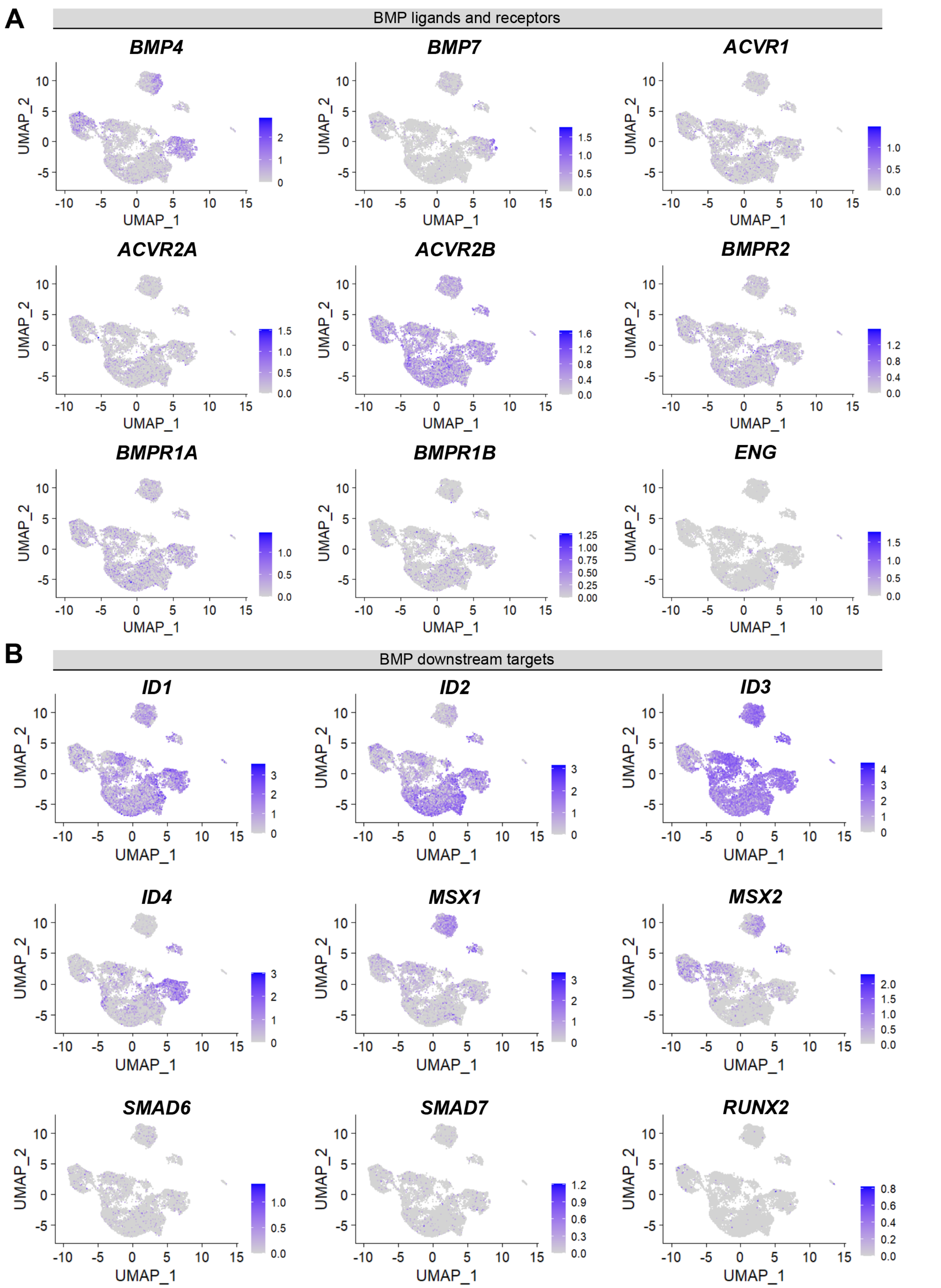


**Figure S5. BMP receptors and downstream targets.**
(**A-B**) UMAP feature plots during D3-9 hiPSC differentiation showing BMP ligands 4 and 7, BMP receptors (**A**) and known BMP downstream targets (**B**).

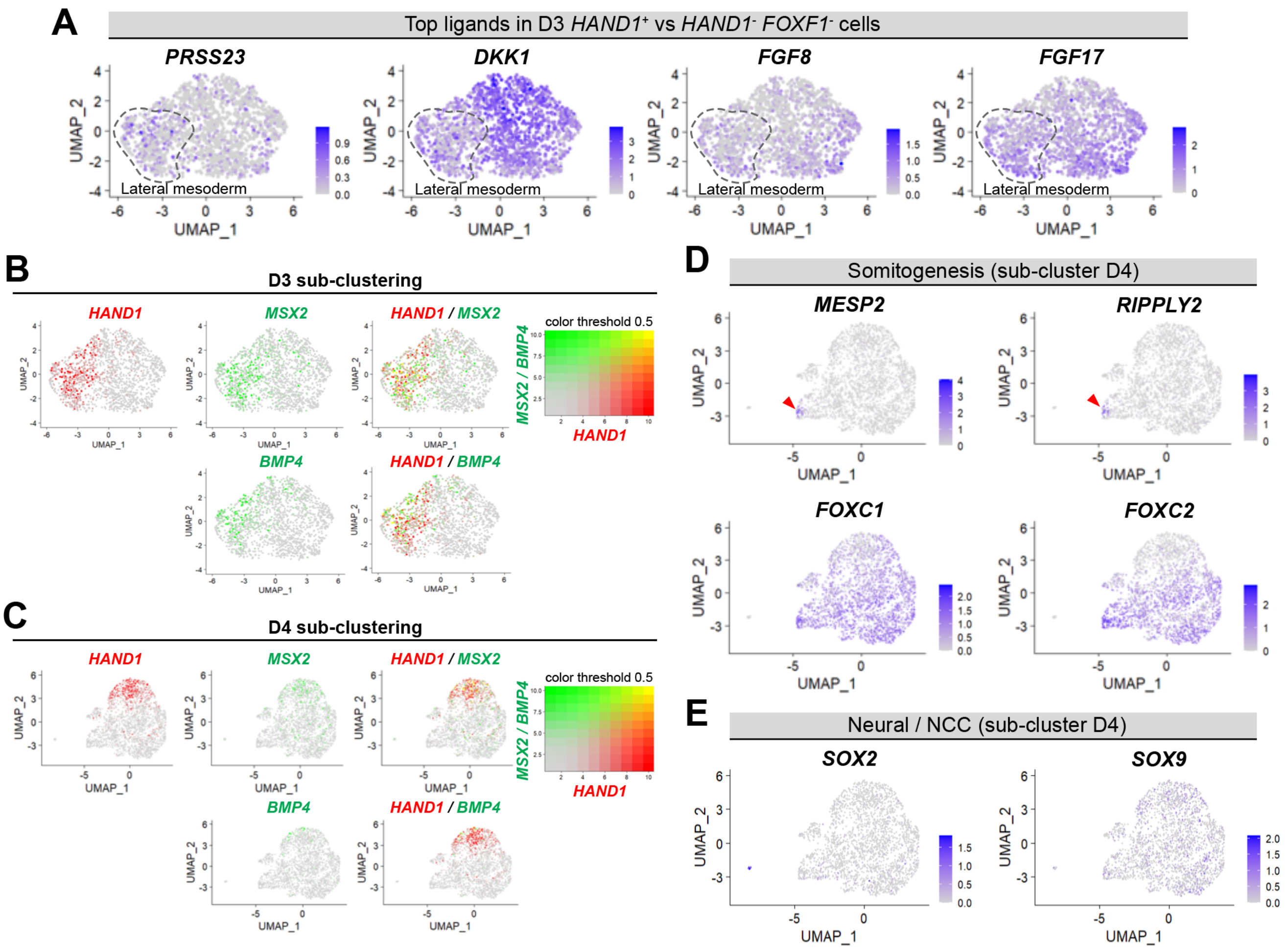


**Figure S6. Sub-clustering of D3 and D4 mesoderm populations.**
(**A**) UMAP feature plots in D3 sub-clustered cells for *PRSS23*, *DKK1*, *FGF8* and *FGF17*. Dashed line shows the approximate location of D3 lateral mesoderm. (**B-C**) UMAP feature plots shows the expression and overlap of *HAND1* versus that of *BMP4* and *MSX2* in D3 (**B**) and D4 (**C**) sub-clustered cells. (**D-E**) UMAP feature plots in D4 sub-clustered cells of markers associated with somitogenesis (**D**) or neural / neural crest cells (NCC, **E**). Red arrowheads indicate few cells expressing somitic markers *MESP2* and *RIPPLY2*.

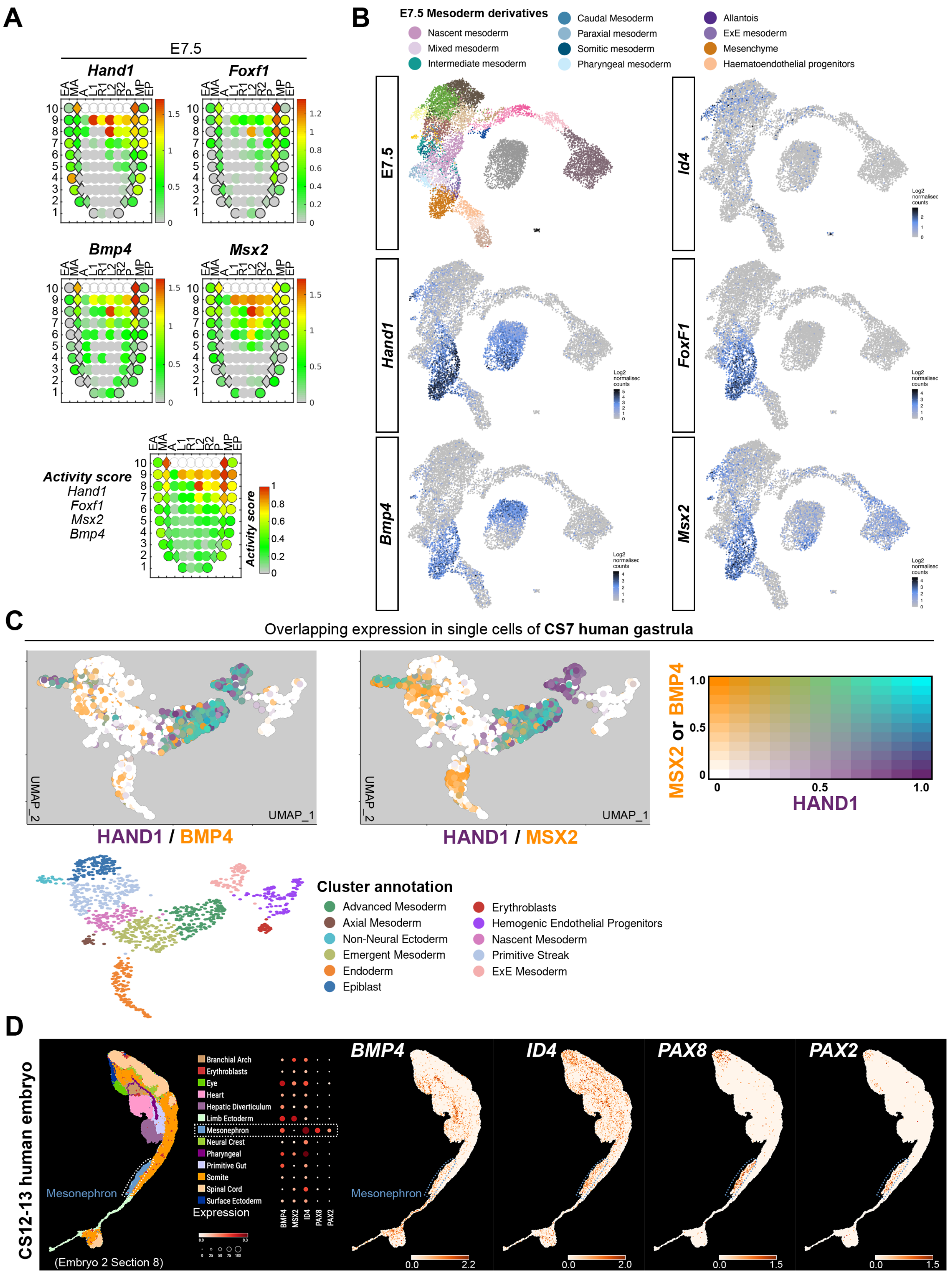


**Figure S7. Human and mouse embryos show different downstream response to BMP4 in LPM and IM tissues.**

**Figure S7. Human and mouse embryos show different downstream response to BMP4 in LPM and IM tissues.**
(**A**) Corn plots generated by the eGastrulation tool[51] shows spatial gene expression and gene activity score calculation at E7.5 for *Foxf1*, *Hand1*, *Bmp4* and *Msx2*. The E7.5 embryo is represented by anatomical tissue slices with an anterior-posterior, left-right orientation along the proximal-distal axis (with 10 being most proximal and 1 most distal). Labels are EA, anterior endoderm; MA, anterior mesoderm; A, anterior epiblast; L1, anterior left lateral; R1, anterior right lateral; L2, posterior left lateral; R2, posterior right lateral; P, posterior epiblast; MP, posterior mesoderm; EP, posterior endoderm. Values are $\log_{10}$ (FPKM+1) transformed for single genes. Normalized activity scores are calculated across E2.5-E7.5 embryo stages and scaled to [0,1]. (**B**) *Foxf1*, *Hand1*, *Bmp4* and *Msx2* and *Id4* expression in single cells of E7.5 mouse embryos, created with the online tool of a mouse gastrulation single cell expression atlas[52]. Cell cluster nomenclature from the source publication was kept and relevant mesoderm clusters are shown. (**C**) Overlapping gene expression of *HAND1* (purple) with *MSX2* or *BMP4* (yellow) in a Carnagie-Stage (CS) 7 human embryo, created with an online tool[20]. Overlapping expression between two genes is shown in the legend with cyan showing the most overlap. (**D**) Spatial transcriptomics through a section of a CS12-13 human embryo shows annotated tissues with different colors, a dot plot graph of *BMP4*, *ID4*, *PAX8* and *PAX2* expression in the same section, and spatial gene expression. The mesonephron lineage is marked by a dotted line. All images were created by the online tool[50].

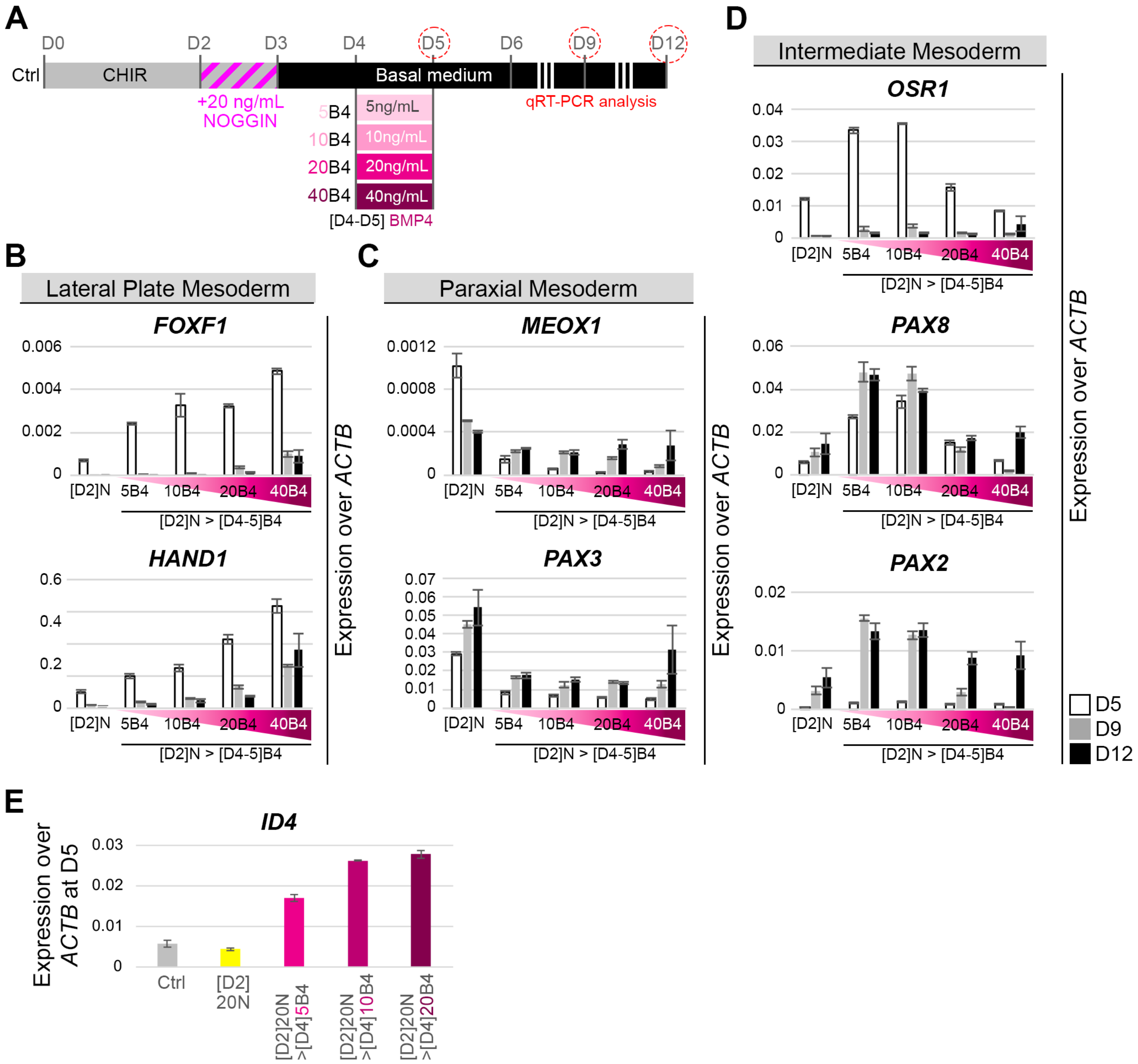


**Figure S8. Early BMP inhibition followed by later BMP4 treatment dose-dependently drives mesodermal lineage allocation.**

(**A**) Schematic showing experiment setup of hiPSC differentiation. Cells were treated with 4 µM CHIR99021 (CHIR) for 72 h and 20 ng/mL NOGGIN from D2-3, followed by basal medium culture until D12 (Ctrl condition). An additional 24 h BMP4 pulse (5, 10, 20, or 40 ng/mL) was given on D4-5. (**B-D**) Cells were collected on D5, D9, and D12 for qRT-PCR analysis for *FOXF1* and *HAND1* (**B**), *MEOX1* and *PAX3* (**C**) or *OSR1*, *PAX8* and *PAX2* (**D**), indicative of LPM, PXM or IM differentiation, respectively. Labels show the indicated treatment conditions shown in A. (**E**) Expression of *ID4* transcript at D5 in different treatments. Ctrl: D0-D3 in 4 µM CHIR, followed by basal medium for two more days (grey). [D2]20N, Ctrl treatment plus 20 ng/mL NOGGIN added at D2 to D3, followed by basal medium (yellow). Pink columns are [D2]20N treatment followed by 24 h BMP4 addition on D4-5, with three concentrations compared: 5, 10 and 20 ng/mL. Column color denotes the predominant fate (grey, control; cyan, LPM; yellow, PXM; pink, IM). Data is from one representative experiment out of three independent experiments with similar outcome.

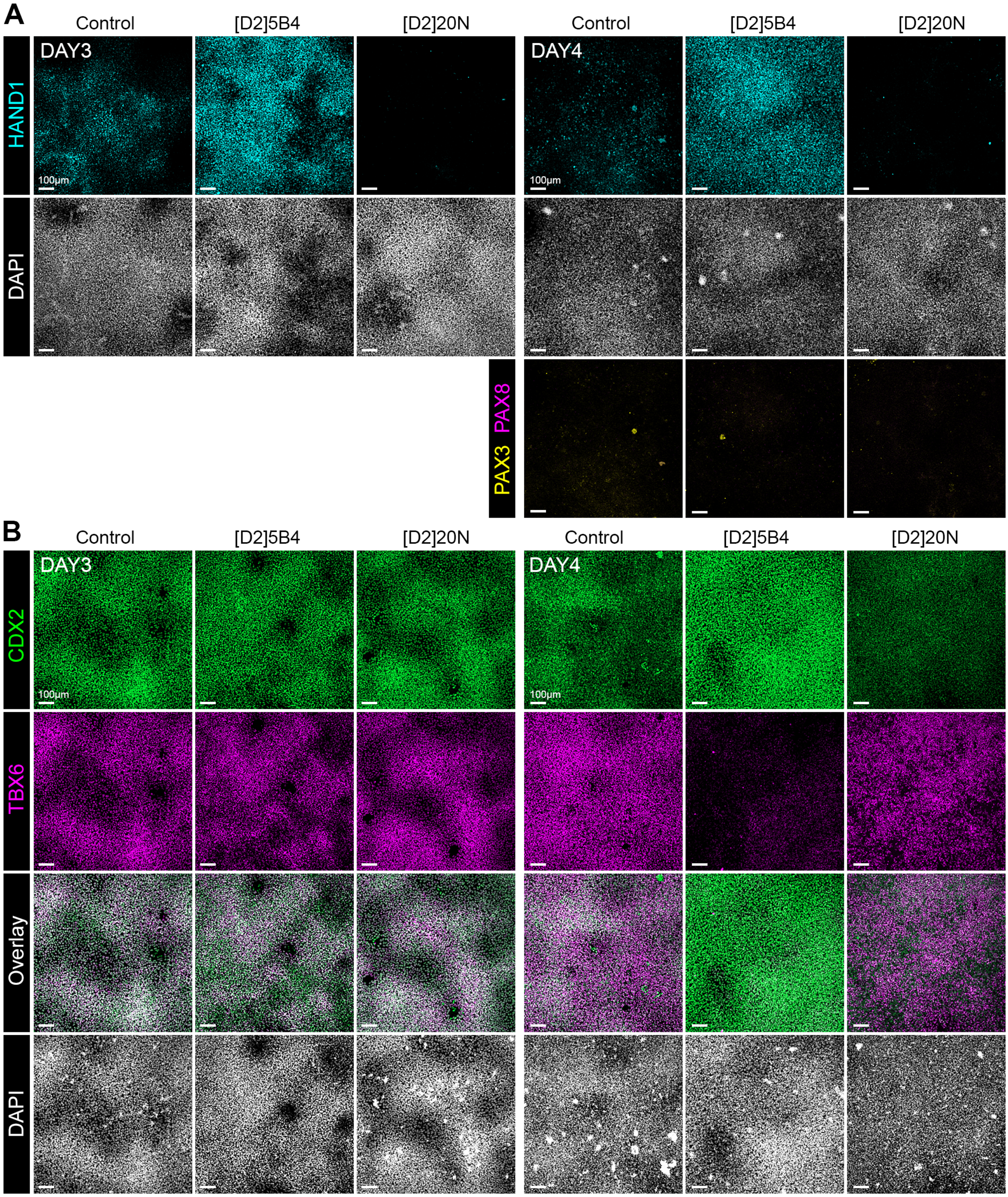


**Figure S9. D5 mesoderm fate changes occur via conversion, not selection.**

(**A**) Left, immunofluorescent staining at D3 for LPM marker HAND1 in 24 h-treated D2 cells with either 5 ng/mL BMP4 or 20 ng/mL NOGGIN ([D2]5B4 and [D2]20N, respectively). Right, immunofluorescent staining at D4 for HAND1, PAX3 and PAX8 in the same populations. (**B**) Immunofluorescent staining for mesoderm progenitor markers CDX2 and TBX6 in the same [D2-3]-treated populations, analyzed at D3 and D4. Nuclear counterstaining was performed with DAPI.

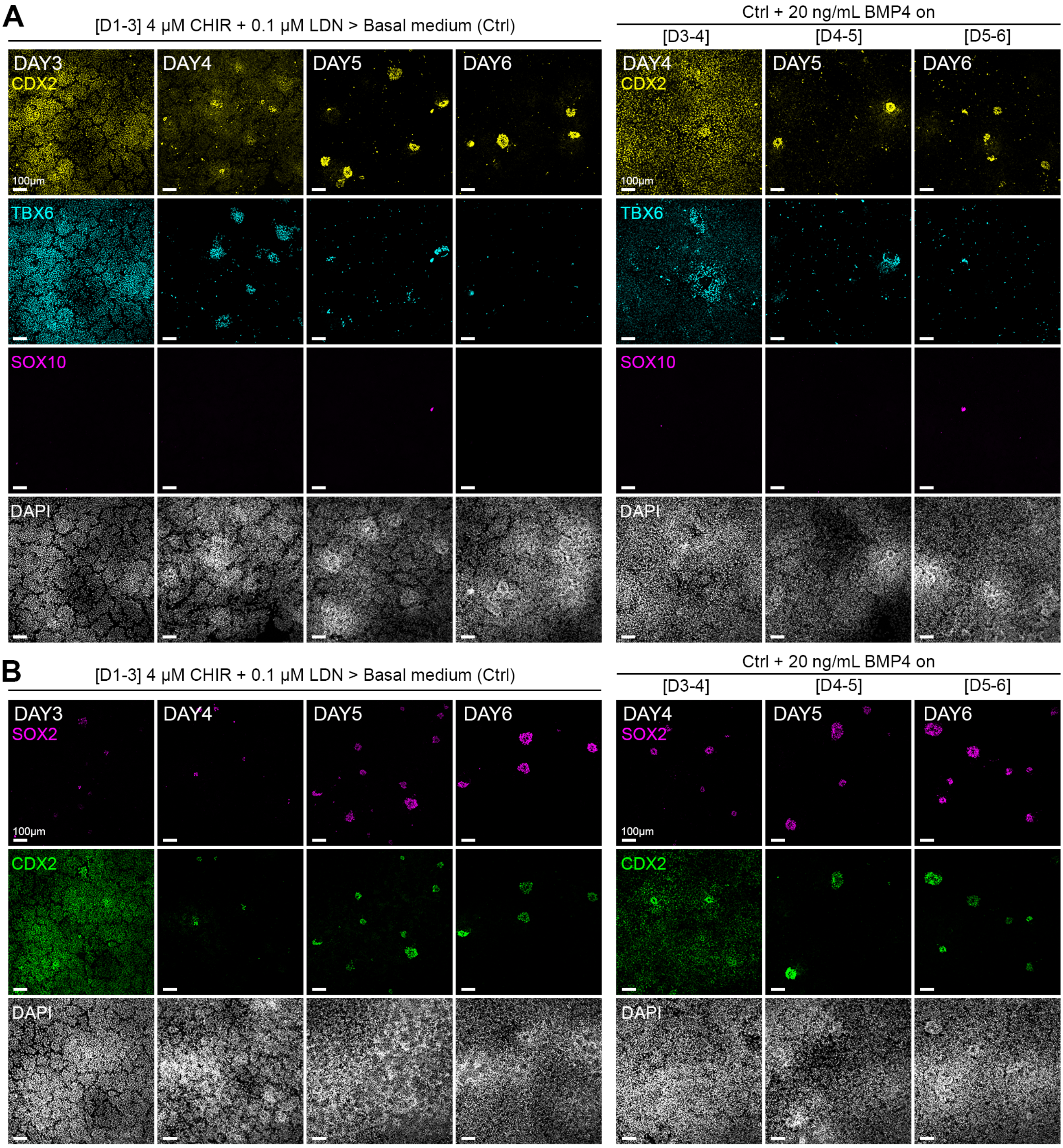


**Figure S10. Spatiotemporal expression of mesoderm progenitor and neural markers during differentiation.**
(**A-B**) Immunofluorescent staining for mesoderm progenitor markers CDX2 and TBX6, and neural crest marker SOX10 (**A**) or CDX2 and neural marker SOX2 (**B**), under various culture conditions. Right, cells stained from D3-6 under the Ctrl protocol (4 µM CHIR99021 and 0.1 µM LDN193189 for 72 h, followed by basal medium culture from D3) or left, Ctrl protocol with additional 20 ng/mL BMP4 added to the culture during 24 h at D3, D4 or D5. Nuclear counterstaining was performed with DAPI. Whereas no SOX10 could be detected, CDX2$^{high}$ cells co-expressed SOX2 in core-like structures. Under additional BMP4 treatment, cells at D4 showed higher TBX6 and CDX2$^{low}$ expression. A lower concentration of 5ng/mL BMP4 was tested with similar results to 20 ng/mL BMP4 (*data not shown*).

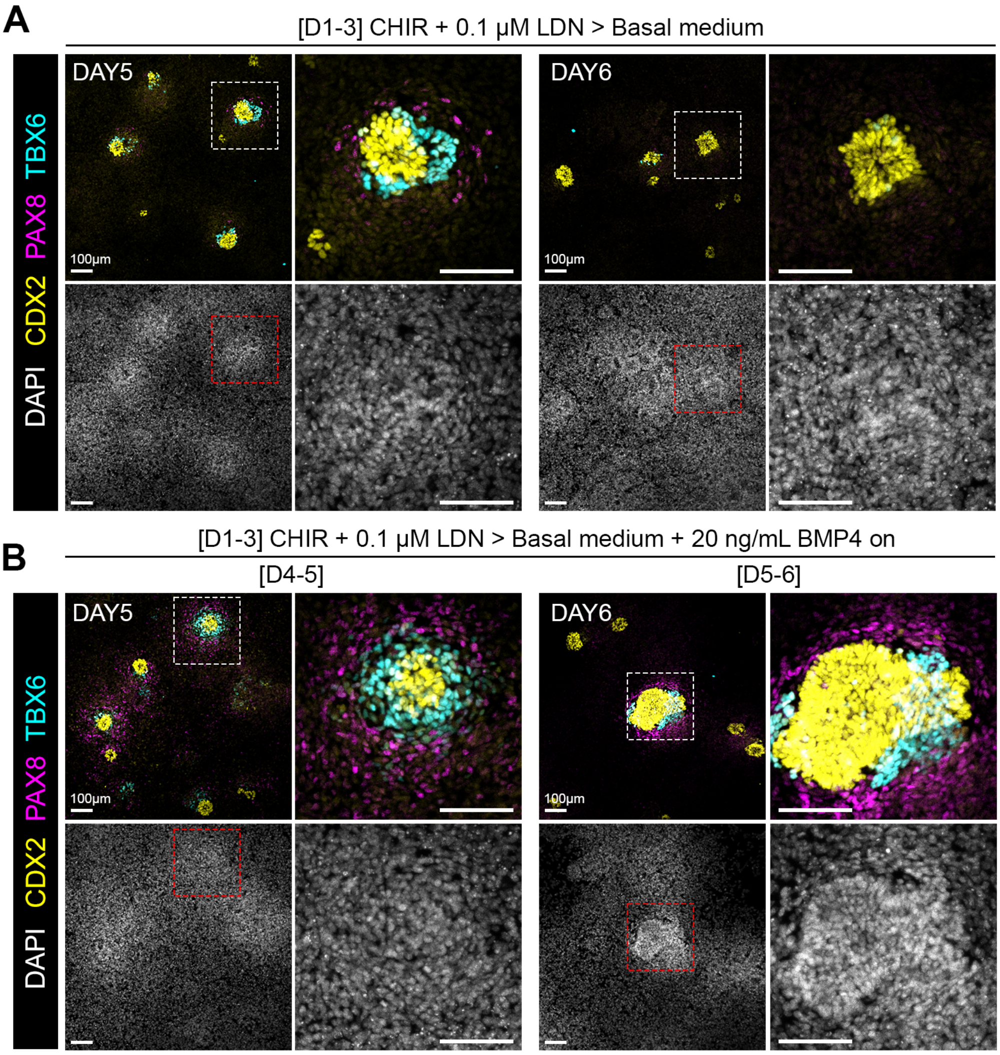


**Figure S11. Spatial organization of IM cells relative to mesoderm progenitor structures.**
(**A-B**) Immunofluorescent staining of mesoderm progenitor markers TBX6 and CDX2, and the IM marker PAX8 on D5 and D6 of differentiation. (**A**) D5 and D6 cells stained under the control protocol (Ctrl): 4 µM CHIR99021 and 0.1 µM LDN193189 for 72 h, followed by basal medium culture from D3. (**B**) D5 and D6 cells stained under the Ctrl protocol with additional 20 ng/mL BMP4 added to the culture during 24 h at D4 or D5. Specific expression patterns were observed in which PAX8$^{+}$ IM cells surrounded TBX6 cells, which in turn surrounded CDX2$^{high}$ cells. A lower concentration of 5ng/mL BMP4 was tested with similar results to 20 ng/mL BMP4 (*data not shown*).
